# Unraveling the Role of P450 Reductase in Herbicide Metabolic Resistance Mechanism

**DOI:** 10.1101/2023.12.19.572429

**Authors:** Carlos Alberto Gonsiorkiewicz Rigon, Satoshi Iwakami, Todd A. Gaines, Franck E. Dayan

## Abstract

Plants require cytochrome P450 reductase (CPR) to supply two electrons for cytochrome P450 monooxygenase enzymes (P450) to react with an organic substrate. The transfer of electrons to the P450 active site in the P450 catalytic site relies on a robust and intricate CPR:P450 complex in the endoplasmic reticulum membrane. Transgenic Arabidopsis plants carrying *CYP81A12* from *Echinochloa phyllopogon*, which metabolizes a broad spectrum of herbicides, were crossed with CPR knockout *atr1* or *atr2* mutant lines. Homozygous gene knockout was confirmed using PCR, and gene copy number of *CYP81A12* was determined using ddPCR. Arabidopsis lines expressing *CYP81A12* in combination with *atr1* or *atr2* knockout were used for herbicide dose-response and metabolism studies. Knocking out *ATR1* in transgenic Arabidopsis *CYP81A12* significantly reduced herbicide resistance. Transgenic mutant plants (*CYP81A12 atr1-b*) had a 3.6-, 5.6-, 6.8- and at least 26-fold reduction in resistance to mesotrione, 2,4-D, penoxsulam and chlorsulfuron, respectively, in the dose-response assay. Knockouts of the *ATR2* also decreased herbicide resistance, but to a lower magnitude than *ATR1*. These results corroborate ½ MS medium assay, and herbicide resistance reduction was observed for additional tested herbicides, bensulfuron, propoxycarbazone and bentazon. Our findings highlight the importance of CPRs in metabolic herbicide resistance in plants, by identifying that a single CPR knockout can reverse herbicide sensitivity . The different CPRs found in weeds have potential as target genes to manage metabolic herbicide resistance evolution. We further provide an in-depth exploration of the evolutionary implications in weed management arising from the results.

**HIGHLIGHTS:** Knocking out cytochrome P450 reductase 1 in herbicide tolerant Arabidopsis reduces herbicide resistance, highlighting CPRs as targets for managing herbicide resistance evolution in weeds.

## INTRODUCTION

Plants are sessile organisms, which remain permanently anchored to their site of germination. To overcome their lack of mobility, plants have evolved complex detoxification mechanisms to deal with environmental and biotic stresses. These natural detoxification processes by chance enable enhanced herbicide metabolic pathways in certain crops, conferring natural tolerance to herbicides (Jeschke *et al*., 2019). This strategic approach has yielded substantial advantages in the commercialization of herbicides intended to eliminate weeds while safeguarding crops; however, the prolonged application of herbicides in agroecosystems has selected weeds that exhibit the same mechanisms of resistance (Powles & Yu, 2010) . Among these, the enhanced capacity of herbicide metabolism is one of the most important (Rigon *et al*., 2020). These selected weeds have an enhanced activity of enzyme families that metabolize the herbicides before reaching the target site (Gaines *et al*., 2020). This mechanism of resistance threatens weed management due to its broad substrate specificity to many herbicides from different mechanisms of action (Rigon *et al*., 2020). Metabolic herbicide resistance mechanism is divided in three phases, known as Phase I – oxidation, Phase II – conjugation, and Phase III – sequestration and degradation. Phase I is mediated primarily by cytochrome P450 monooxygenase (P450) enzymes. These enzymes are membrane-bound, localized in the endoplasmic reticulum, and are involved in the biosynthesis of many molecules including antioxidants, secondary metabolites, and in detoxification of many xenobiotic compounds (Pandian *et al*., 2020). P450s are the most important enzymes in phase I of herbicide metabolism and plants possess a high number of P450 genes that might play a crucial role in herbicide resistance in metabolic resistant weed species (Dimaano & Iwakami, 2021).

Cytochrome P450 reductase (CPR) is a membrane-bound enzyme localized in the endoplasmic reticulum that plays a critical role in the biosynthesis of a wide range of compounds in living organisms, including plants, animals, and bacteria. The structure of P450 reductase is divided in four domains, including *N*-terminal flavin mononucleotide (FMN)-binding domain, a connecting domain, an flavin adenine dinucleotide (FAD)-binding domain, and a *C*-terminal NADPH-binding domain (Wang *et al*., 1997, Jensen & Møller, 2010) (Fig. **1**). CPR enzymes arose from of a fusion of different genes encoding FAD-binding, ferredoxin-NADP^+^, and an FMN-flavodoxin (Porter & Kasper, 1986). Plants require CPR to supply two electrons for P450 enzymes to conduct each monooxygenase reaction with an organic substrate (Lau & O’Keefe, 1996, Paine *et al*., 2005). The CPR reduction includes multiple steps for electron transfer. The first step is reduction from NADPH to FAD, then FMN, and finally to the P450 acceptor (Munro *et al*., 2001) (Fig. **1**). The transfer of electrons to the P450 active site in the P450 catalytic cycle relies on a robust and intricate CPR:P450 complex in the membrane (Mukherjee *et al*., 2021), making CPR function an essential component of specialized metabolism.

**Fig. 1.**
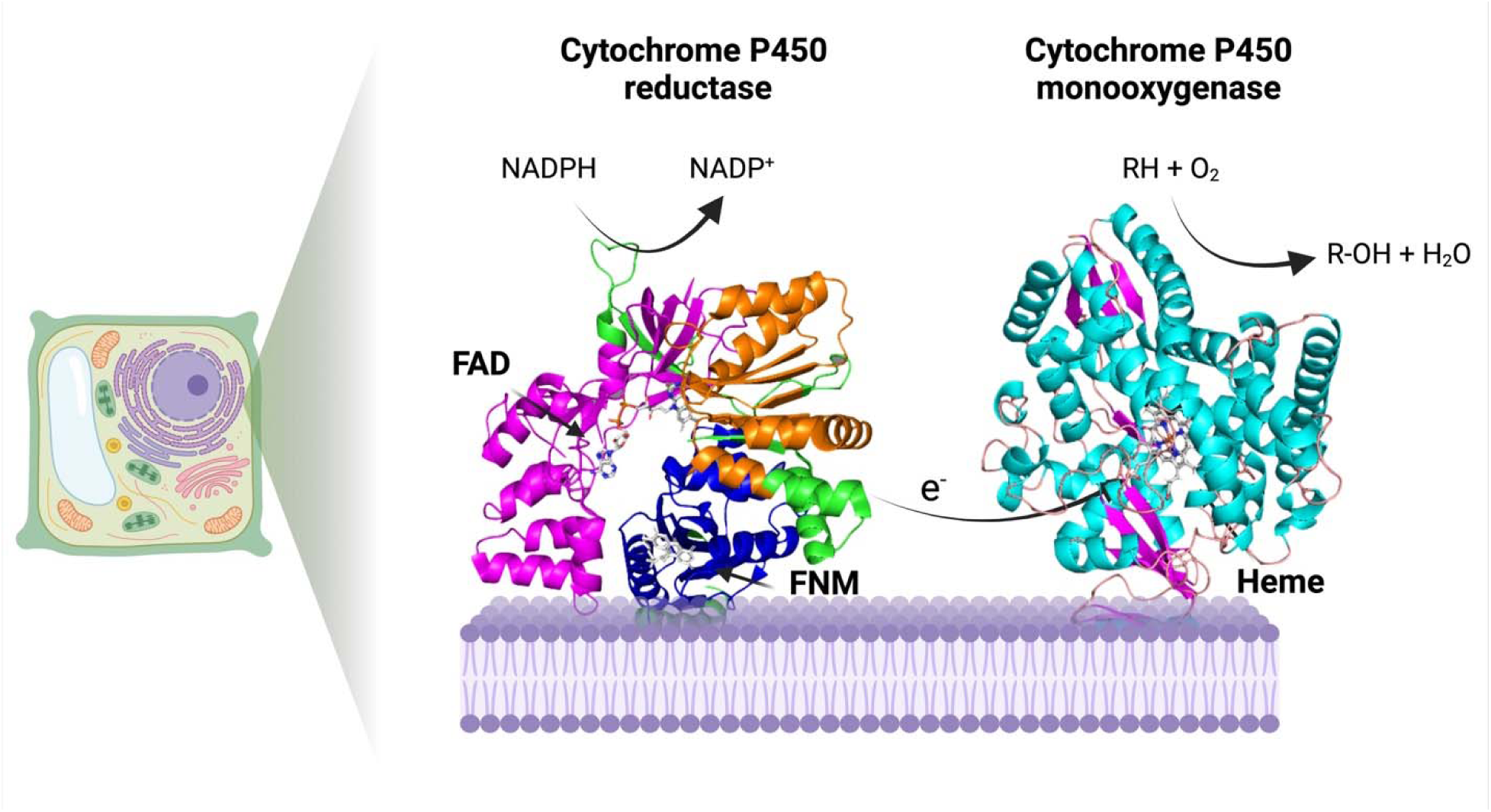
Process of electron transfer from NADPH-cytochrome P450 reductase to the catalytic site of cytochrome P450 monooxygenase. Two electrons are sequentially transferred from NADPH to the heme group of P450 through the cofactors FAD and FMN, each transferred in single-electron steps. Cytochrome P450, once activated by the electron transfer, catalyzes the oxidative breakdown of a substrate by inserting one oxygen atom into the chemical compound. Created with Biorender.

**Fig. 2.**
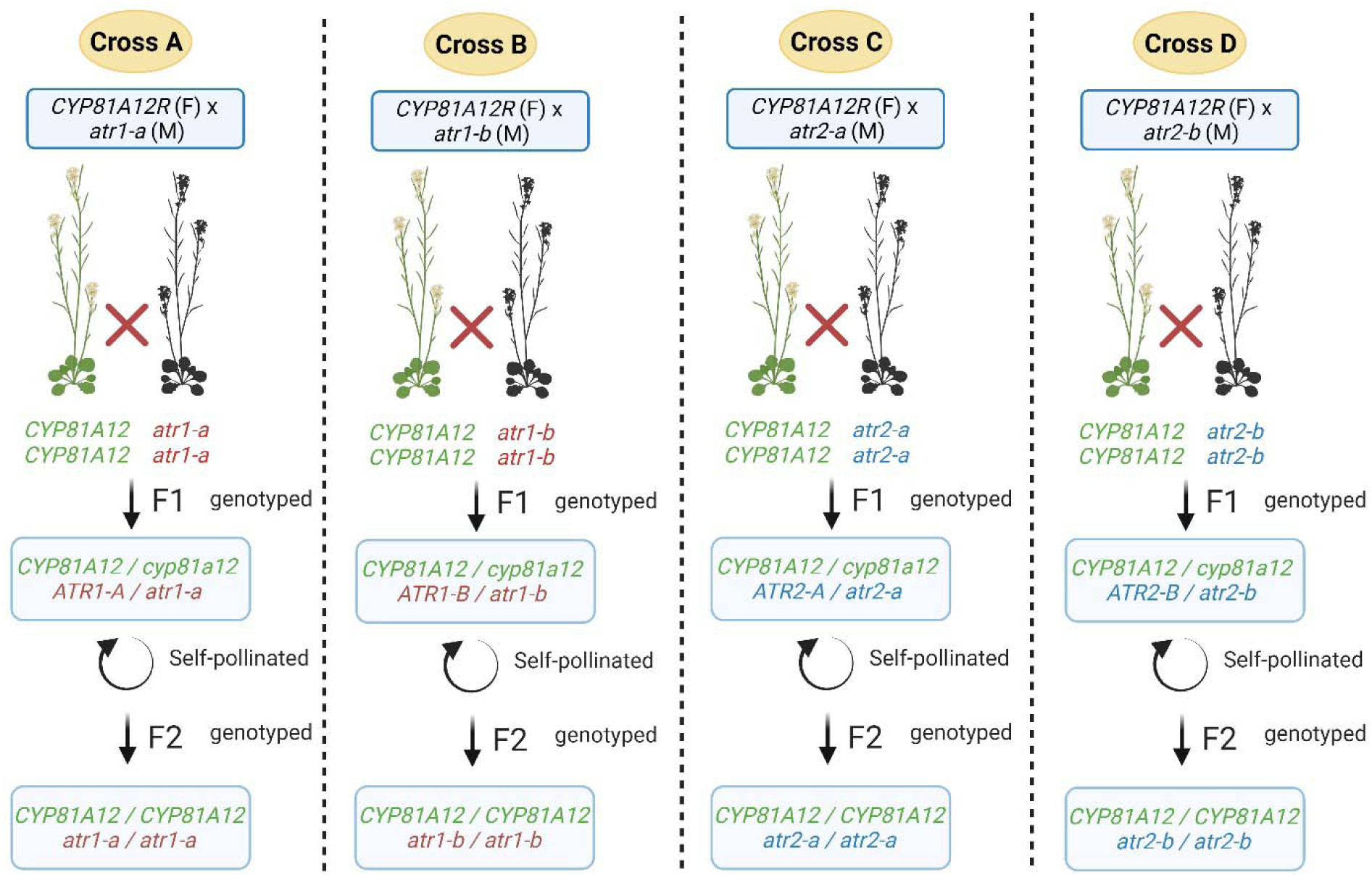
Schemes representing the Arabidopsis crosses to generate transgenic Arabidopsis *CYP81A12* with loss of function in *atr1* or *atr2*. F1 plants were genotyped for T-DNA insertion and *CYP81A12* presence. F2 plants were genotyped for homozygous T-DNA insertion and two copies of *CYP81A12* using ddPCR. M - male, F - female. Created with BioRender.com.

In contrast to the huge number of P450 genes within plant genomes, for example, 246 in *Arabidopsis thaliana* (Arabidopsis) (Paquette *et al*., 2009), 328 in *Oryza* spp. (rice) (Nelson *et al*., 2004), 867 in *Echinochloa crus-galli* (barnyardgrass) (Wu *et al*., 2022), 694 in *E. colona* (jungle rice) (Wu *et al*., 2022), and 323 in *Conyza canadensis* (horseweed) (Peng *et al*., 2014), CPRs generally have only two or three copies in plant genomes. There are two CPR genes in Arabidopsis, referred to as *Arabidopsis thaliana* cytochrome P450 reductase 1 (*ATR1*), 2 (*ATR2*) (Urban *et al*., 1997) and one hypothetical reductase 3 (*ATR3)* that encodes a diflavin reductase essential for embryogenesis, but that is unable to reduce P450 *in vitro* (Mizutani & Ohta, 1998, Varadarajan *et al*., 2010). The isoforms are grouped in two distinct groups with different roles. Class I is *ATR1*, which is involved in primary and basal metabolism. Class II is *ATR2*, which is inducible by environmental stimuli such as wounding, pathogen attack, or high light intensity exposure (Rana *et al*., 2013, Parage *et al*., 2016).

In current research efforts, there has been a growing focus on the identification of P450 enzymes responsible for herbicide metabolism in weeds (Rigon *et al*., 2020). An important example is the CYP81A12 enzyme derived from a multiple-resistant population of *Echinochloa phyllopogon*. This enzyme is capable of metabolizing herbicides from diverse modes of action when expressed heterologously in *E. coli* or Arabidopsis (Iwakami *et al*., 2014, Dimaano *et al*., 2020). Considering the dependence of P450 enzymes on P450 reductase for their functionality, the validation of this interdependence and the identification of pivotal cytochrome P450 reductases could potentially offer a valuable mechanism for weed management and the mitigation of metabolic herbicide resistance evolution in weeds.

The study hypothesized that *ATR1* and *ATR2* are the primary electron donors for P450 enzymes and that silencing these genes would reduce the activity of P450s involved in herbicide metabolism. The main objective was to assess the potential of targeting cytochrome P450 reductase to diminish herbicide metabolism mediated by *CYP81A12*.

## MATERIALS AND METHODS

### Plant material

Mutant SALK lines of Arabidopsis were ordered from the Arabidopsis Biological Resource Center (ABRC - https://abrc.osu.edu/). The lines SALK_111558, SALK_208483C have a T-DNA insertion in Arabidopsis cytochrome P450 reductase 1 (*ATR1* - at4g24520), hereafter named as *atr1-a* and *atr1-b*, respectively. The lines SALK_026053 and SALK_152766 have T-DNA insertion in Arabidopsis cytochrome P450 reductase 2 (*ATR2* - at4g30210), hereafter named as *atr2-a* and *atr2-b*.

Seeds from mutants and wild-type Columbia_0 (Col_0) were sown in soil and kept in a growth chamber at 22/18°C with a light intensity of 120 µE m^-2^ s^-1^ for 16 h. When seedlings had four true leaves, they were transplanted into individual pots and kept in the growth chamber in the same conditions. All the following experiments used the same methods to grow the Arabidopsis plants.

### Characterization and confirmation of T-DNA insertion

PCR assays were performed to identify and confirm the presence of T-DNA insertion in the *ATR1* and *ATR2* genes in the mutant lines and to verify homozygosity. The methods are described in Methods **S1**.

### Generating transgenic herbicide-resistant Arabidopsis with mutant *atr1* or *atr2* genes and double knockout lines

Aiming to generate transgenic Arabidopsis lines with a loss of function in *ATR1* or *ATR2*, a transgenic Arabidopsis line carrying the P450 *CYP81A12* from *Echinochloa phyllopogon* (synonym *E. oryzicola*, late watergrass) (Iwakami *et al*., 2014) was used. The transgenic Arabidopsis metabolizes five different herbicide modes of action including acetolactate synthase (ALS) inhibitors (bensulfuron-methyl, pyrazosulfuron-ethyl, chlorsulfuron, azimsulfuron, imazamox, propoxycarbazone-sodium and penoxsulam), 4-hydroxyphenylpyruvate dioxygenase (HPPD) inhibitor (mesotrione), protoporphyrinogen oxidase (PPO) inhibitor (pyraclonil), deoxyxylulose phosphate synthase (DXS) inhibitor (clomazone), and photosystem II (PSII) inhibitor (bentazon) (Dimaano *et al*., 2020). This population hereafter is called Arabidopsis *CYP81A12*. This line was crossed with mutant lines *atr1-a*, *atr1-b*, *atr2-a*, and *atr2-b* to generate transgenic CYP81A12 carrying mutant *atr1* or *atr2*, hereafter are named as *CYP81A12 atr1-a*, *CYP81A12 atr1-b*, *CYP81A12 atr2-a*, and *CYP81A12 atr2-b*. The methods of crossing and genotyping populations are described in Methods **S2**. Aiming to generate double knockout lines, i.e. same line with loss of function in *atr1* and *atr2*, mutants plants were crossed and the methods are described in Methods **S3**.

### Gene expression analysis in *atr1* and *atr2* mutant lines

Mutant and transgenic mutant lines were grown in a growth chamber with the same conditions as specified before. Approximately 100 mg of leaf tissue were collected from plants in the rosette stage. Sampling consisted of cutting the youngest fully expanded leaf for each biological replication, placing in 2 mL tubes, and incubating immediately in liquid nitrogen and afterwards kept at -80°C. Plants of lines Col_0, transgenic *CYP81A12*, and transgenic mutants *CYP81A12 atr1-a*, *CYP81A12 atr1-b*, *CYP81A12 atr2-a*, and *CYP81A12 atr2-b* were kept for fresh mass measurement at 28 d after transplant to analyze any fitness cost in the mutant lines.

RNA extraction was conducted using Direct-zol™ RNA miniprep from Zymo Research following manufacturer instructions. DNase I treatment was conducted during the RNA extraction protocol. Quantification and quality control were performed using Nanodrop 2000. cDNA synthesis was conducted using iScript™ from Zymo Research using 1 µg of RNA. Gene expression was analyzed on CFX96 Real Time System using SsoAdvanced™ Universal SYBR® Green Supermix. The reactions consisted of 5 μL of cDNA from samples diluted in Nuclease-Free Water in 1:20, 2.5 μL of each primer at 5 μM, and 5 μL of SsoAdvanced universal SYBR® Green supermix (2X). Thermal cycler conditions consisted of an initial cycle of 30 s at 95°C, followed by a sequence of 35 cycles starting at 95°C for 10 s and 72°C for 30 s. Melt-curve analysis was performed by adding steps incrementing temperature 65–95°C by 0.5°C.

Primer sets were specifically designed to target various regions of the genes, both upstream and downstream of the T-DNA insertion site. This systematic design allowed for accurate analysis of how the gene was affected by the T-DNA insertion. For *ATR1* gene, four sets of primers were designed, called ATR1_A with forward primers in the 5’UTR and reverse primers in the 1^st^ exon (spanning the T-DNA insertion position in *atr1-a* line), ATR1_B with forward and reverse primers in the 1^st^ exon, ATR1_C with the forward primer in exon 4^th^ and reverse primer in exon 5^th^ (spanning the T-DNA insertion position in *atr1-b* line), and ATR1_D with forward in the 8^th^ exon and reverse in the 9^th^ exon (Fig. **3**). For *ATR2* gene, five sets of primers were designed, ATR2_A with forward and reverse primers localized in the 1^st^ exon, ATR2_B with forward primer in the 3^th^ exon and reverse primer in the 4^th^ exon (spanning the T-DNA insertion position in *atr2-a* line), ATR2_C with forward primer localized in the 9^th^ and reverse in the 10^th^ exon, ATR2_D with forward primer in the 12^th^ exon and reverse in the 13^th^ (spanning the T-DNA insertion position in *atr2-b* line), and ATR2_E with forward and reverse primers designed in the 14^th^ exon. The primer locations in *ATR1* and *ATR2* are represented in Fig. **3a** and **b**, respectively. Primer sequences for gene expression analysis and their characteristics are listed in Tab. **S2**.

**Fig. 3.**
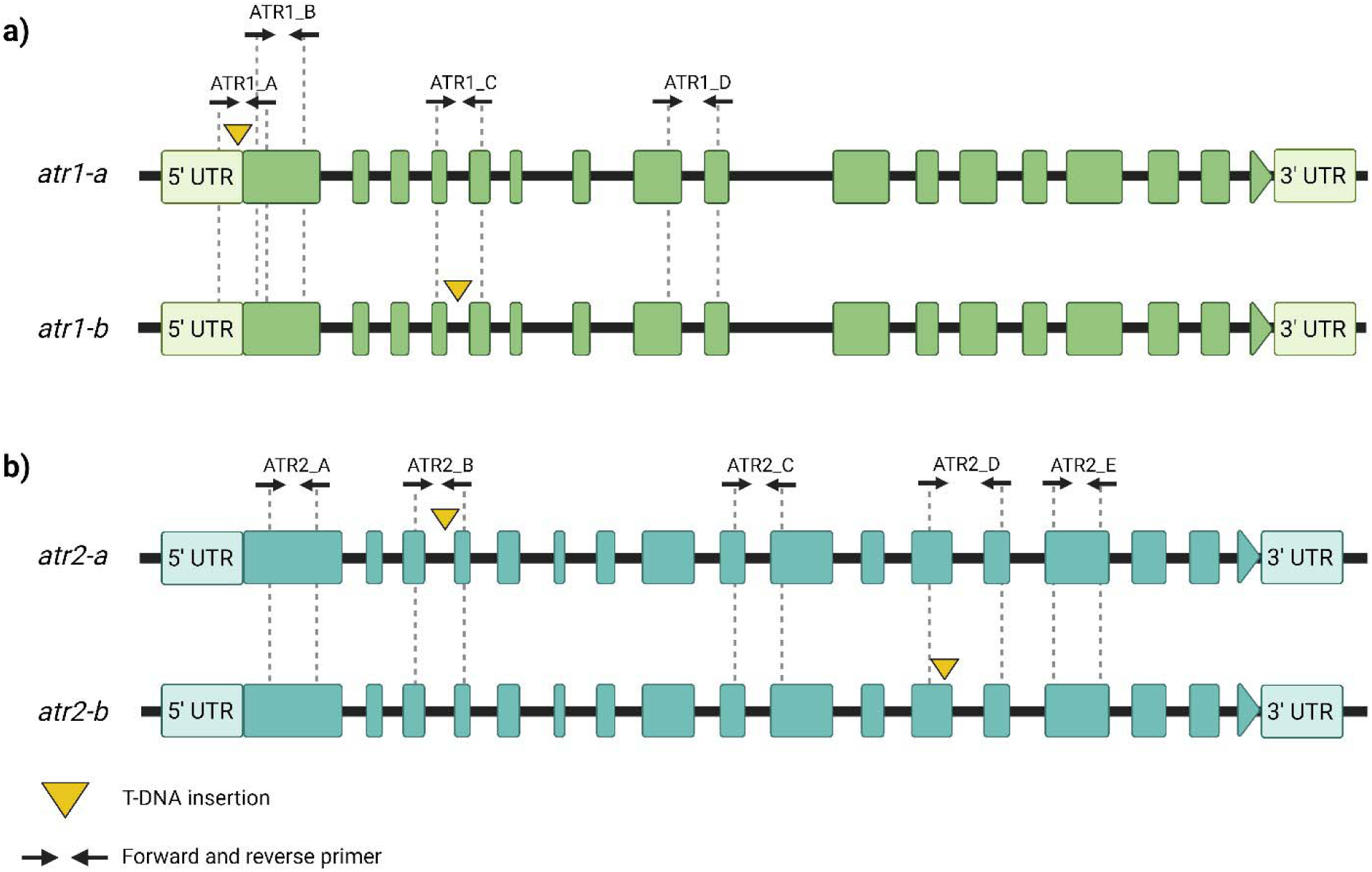
T-DNA location in *ATR1* (A) and *ATR2* (B) genes in SALK lines and primers localization for real time qPCR gene expression analysis. Yellow triangle locates the T-DNA insertion in the genes, *atr1-a* and *atr2-b* have T-DNA insertion in the 5’UTR and 4^th^ intron, respectively, and *atr2-a* and *atr2-b* have T-DNA insertion in the 3^rd^ and 12^th^ exon, respectively. Boxes indicate exons or untranslated regions (UTR), black line indicates introns. Created with BioRender.com.

### Herbicide sensitivity of Arabidopsis lines in ½ MS medium

Plant growth of different lines of Arabidopsis were evaluated on ½ Murashige and Skoog (MS) solid medium containing different doses of bensulfuron, chlorsulfuron, penoxsulam, propoxycarbazone, mesotrione and bentazon. Seeds of Arabidopsis wild type (Col_0), transgenic Arabidopsis *CYP81A12*, mutants and transgenic mutant *CYP81A12 atr1-a*, *CYP81A12 atr1-b*, *CYP81A12 atr2-a*, and *CYP81A12 atr2-b* were sterilized by chloride gas for 7 h by mixing 100 mL of bleach with 3% hydrochloric acid. Technical grade herbicides were diluted in ethanol (bensulfuron, penoxsulam, propoxycarbazone, bentazon) or methanol (chlorsulfuron, mesotrione) to mix with ½ MS medium before it solidified. Petri dishes containing ½ MS with different herbicides doses were divided into 10 sections, where six seeds of each Arabidopsis line were placed. The plates were incubated in a growth chamber at 25°C, 14-h photoperiod, and 110 μE m^-2^ s^-1^ light intensity until the day of evaluation. The number of green leaves were counted 14 d after plating, and the data were normalized to percentage of untreated control.

### Whole-plant dose response

Arabidopsis plants of Col_0, transgenic *CYP81A12*, and transgenic mutant *CYP81A12 atr1-a*, *CYP81A12 atr1-b*, *CYP81A12 atr2-a*, and *CYP81A12 atr2-b* were grown with the same conditions as specified before. Plants in rosette stage (14 d after transplanting) were sprayed with increasing doses of chlorsulfuron (Telar ® XP, 750 g Kg ^-1^), penoxsulam (Ricer, 217 g a.i. kg^-1^), mesotrione (Callisto 480g a.i. L^-1^), and 2,4-D (Shredder 2,4-D LV4, 455g a.e. L^-1^). The herbicides were applied in an automated spray chamber (Greenhouse Spray Chamber, model Generation IV) using a TJ8002E nozzle, calibrated to deliver 200 L ha^-1^ at a pressure of 280 kPa and speed of 1.2 m s^-1^. The experimental design was completely randomized in 6 X 11 with four replications. Factor A was Arabidopsis lines. Factor B was increasing herbicides dose for each herbicide. Chlorsulfuron and penoxsulam doses were in range from 0 to 51.2 g a.i. ha^-1^; mesotrione doses were from 0 to 400 g a.i. ha^-1^; and 2,4-D doses were from 0 to 128 g a.e. ha^-1^. The shoot fresh mass was measured at 28 d after herbicide treatment.

### Herbicide metabolism analysis

Seedlings of Arabidopsis lines Col_0, transgenic *CYP81A12*, and transgenic mutant *CYP81A12 atr-a*, *CYP81A12 atr-b*, *CYP81A12 atr2-a*, and *CYP81A12 atr2-b* were transplanted into pots filled with potting soil and grown under the same conditions as previously described. At rosette stage, mesotrione or chlorsulfuron were applied were applied at 100 and 50 g ha^-1^, respectively. The doses were chosen based on the whole-plant dose-response curve experiment described above. After herbicide application the plants were kept in the growth chamber at same conditions and whole plants were collected at 6, 12, 24, 48, and 96 h for mesotrione and adding 144 h and 196 h for plants treated with chlorsulfuron. At each time of tissue collection, plant samples were weighed, washed with 80% acetone three times, dried with towel paper, placed in 15 mL Falcon tubes, and kept at -80°C ultra-freezer until the extraction. The extraction was conducted by grinding the tissue with a pestle, then added 10X of plant mass with 80% methanol and 0.1% formic acid. The solution was homogenized for 30 s using PowerGen 125 (Fisher Scientific) and centrifuged for 10 min at 4,700Xg (Sorvall Legend X1R, Thermo Scientific, Waltham, MA). The supernatant was filtered with Econofltr Nylon 13 mm 0.2 µm (Agilent Technologies, Santa Clara, CA) and injected in the LC-MS/MS.

LC-MS/MS system consisted of a Nexera X2 UPLC with 2 LC-30AD pumps, a SIL-30AC MP autosampler, a DGU-20A5 Prominence degasser, a CTO-30A column oven, and SPD-M30A diode array detector coupled to an 8040 quadrupole mass-spectrometer (Shimadzu Scientific Instruments, Columbia, MD). For mesotrione, the MS was in positive mode [M+H]^+^ with MRM optimized for mesotrione 340>227.95 and set for 100 ms dwell time with a Q1 pre-bias of - 16.0V, a collision energy of -18.0V and a Q3 pre-bias of -16.0V and for hydroxy-mesotrione 356>55>228 and set for 100 ms dwell time with a Q1 pre-bias of -16V, a collision energy of - 18.0V and a Q3 pre-bias of -16.0V. For chlorsulfuron, the MS was in positive mode with MRM optimized for chlorsulfuron 357.9>141.1>167.05 and set for 100 ms dwell time with a Q1 pre-bias of -25.0V, a collision energy of -20.0V and a Q3 pre-bias of -15.0V and for hydroxy-chlorsulfuron 374.0>127.0>182.0 and set for a collision energy of -35.0. The samples were chromatographed on a 100X4.6 mm Kinetex 2.6 µm biphenyl column (Phenomenex, Torrance, CA) maintained at 40°C. For both herbicides, solvent A consisted of water with 0.1% formic acid and solvent B was methanol with 0.1% formic acid. The solvent program started at 50% B and increased to 95% B in 5 min and maintained at 95% for 4 min. The solvent was returned to 50% B and maintained there for 3 min before the next injection. The flow rate was set at 0.4 mL min^-1^ and each sample were analyzed with 5 µL injection volumes.

### Identification and phylogenetic analysis of CPRs in different weeds

The presence of CPR genes and copy number was searched in several important weed species. The genomes of *Alopecurus myosuroides* (Cai *et al*., 2023), *Chenopodium formosanum* (Jarvis *et al., 2022)*, *Bassia scoparia* (Hall *et al*., 2023), *Eleusine indica* (Zhang *et al*., 2023), *Bromus tectorum* (Revolinski *et al.,* 2023), *Echinochloa crus-galli* (Wu *et al*., 2022), *E. oryzicola* (Wu *et al.,* 2022), *E. colona* (Wu *et al*., 2022), *Ipomoea purpurea* (Gupta *et al*., 2023) *, Poa annua* (Robbins *et al*., 2023), and *Conyza canadensis* (Laforest *et al*., 2020) were accessible through WeedPedia (https://weedpedia.weedgenomics.org) (Montgomery *et al*., 2023) . CPRs were identified with a blast search using *ATR1* (AT4G24520) or *ATR2* (AT4G30210) against the genomes or utilized the InterPro ID for NADPH-cytochrome P450 reductase (IPR023208). Additionally, the genomes of *Amaranthus palmeri* and *A. tuberculatus* were available (Montgomery *et al*., 2020), and the COGE web browser (https://genomevolution.org/coge) was used for blasting purposes. The *CPR* genes in *Amaranthus* species were identified using a comparative analysis approach. Firstly, *ATR1* and *ATR2* sequences were subjected to a sequence similarity search against the *Amaranthus hypochondriacus* (grain amaranth) genome (Lightfoot *et al*., 2017), which is accessible through the Phytozome database. Subsequently, the identified sequences were further compared against the Palmer amaranth and common waterhemp genomes. Other already known plant *CPR* genes were extracted from GeneBank NCBI (Benson *et al*., 2013), such as corn (*Zea mays*, CPR2a: NP_001159331.1, CPR2b: NP_001146345.2), rice (*Oryza sativa*, CPR2a: XP_015651232.1, CPR2b: XP_015650780.1), soybean (*Glycine max*, CPR1: NP_001236742.2, CPR2: XP_003549436.1), cotton (*Gossypium hirsutum*, CPR1: ACN54323.1, CPR2: NP_001314398.2), and Barrel medic (*Medicago truncatula*, CPR1: XP_003602898.1, CPR2: XP_003610109.1). Multiple protein alignment was performed using ClustalO. Tree construction with Neighbor-joining method was performed using Geneious Prime® 2023.0.1. The *N*-terminal, transmembrane, FMN binding, FAD binding, and NADP^+^ binding domains were identified. The transmembrane was identified using DeepTMHMM tool (Hallgren *et al*., 2022) (https://dtu.biolib.com/DeepTMHMM).

### Statistical analysis

Gene expression analysis of *ATR1* and *ATR2* genes in mutant plants was performed using *ALS* as a normalization gene (Chen *et al*., 2017). Expression analyses were performed utilizing four biological and two technical replicates for each line. The mean Ct values and the standard deviation were calculated by treatment. A melt curve analysis confirmed the presence of a single amplified product for each reaction. Relative transcript abundance was calculated using 2^(-ΔΔCt)^ method for each gene and biotype (Livak & Schmittgen, 2001). The reference population used for the calculation was Col_0. Dunnett’s test (p < 0.05) was used to compare the relative expression between Col_0 and mutant lines.

Number of green leaves at 14 d after seeds were planted and shoot fresh weight (% of untreated control) at 28 d after herbicide application (DAA) were used as response variable for the ½ MS medium and whole-plant dose-response experiment, respectively. Data were converted into percentages relative to untreated treatment. The drc package (Ritz *et al*., 2015) from statistical software R v.3.5.3 (R Core Team, 2023) was used for data analysis. The data were tested for the best nonlinear model by comparing different models by *ANOVA()* and using function *modelFit()* testing for lack-of-fit. Based on the analysis, the data were adjusted using the three-parameter log-logistic model with the function *modelFit()*. For the whole-plant dose response experiment, the herbicide doses causing 50% reduction in shoot fresh weight (GR_50_) were determined using the function *summary()* and the resistance index (RI) was calculated by the function *EDcomp()* by dividing the GR_50_ from transgenic *CYP81A12* or transgenic *CYP81A12* mutants (*atr1* or *atr2*) by the GR_50_ of Col_0. For the dose-response in ½ MS medium the calculation was the same, using the GR_50_ obtained by the variable green leaves. The *x* and *y* data from the dose-response curves of both experiments were extracted from R using the function *write.csv()* and plotted with the treatment means by GraphPad Prism version 9 for Windows (San Diego, California).

The metabolism data obtained by LC-MS/MS were submitted to analysis of variance (ANOVA) and the means were compared using Dunnett’s test (p < 0.05). The peak area of the herbicides and metabolites of all the different Arabidopsis lines were compared to the Col_0 line.

## RESULTS

### Arabidopsis mutants

SALK lines *atr1-a*, *atr1-b*, and *atr2-a* were segregating for T-DNA insertion and heterozygous plants were selected to self and seed kept for seed production. *atr2-b* was homozygous for T-DNA insertion and four plants were self-pollinated to produce seeds. Progeny plants from *atr1-a*, *atr1-b*, and *atr2-a* were genotyped again and individuals homozygous for the T-DNA insertion were selected to self and seeds were produced. Homozygous T-DNA insertion in *ATR1* in *atr1-a* and *atr1-b* lines and homozygous T-DNA insertion in *ATR2* in *atr2-a* and *atr2-b* lines were confirmed by PCR. Primers LP and RP for both genes amplified only when Col_0 DNA was in the reaction. No SALK line amplified the WT version of the respective gene and only amplified the T-DNA reaction, confirming the homozygosity for T-DNA insertion. The amplifications were confirmed by electrophoresis in 1% agarose gel (Fig. **S3**), followed by Sanger sequencing. Based on Sanger sequencing results, the insertion of T-DNA was in the 5’ UTR and in the 4^th^ intron in *atr1-a and atr1-b* mutant lines, respectively, for *ATR1,* and in the 4^th^ intron and the 12^th^ exon in *atr2-a* and *atr2-b* mutant lines, respectively, for *ATR2* (Fig. **S3**).

### Transgenic Arabidopsis *CYP81A12* with *atr1* and *atr2* knockdown

F1 plants from crosses were genotyped for T-DNA insertion. Around 20 plants were genotyped from each cross and around 90% of the plants were heterozygous for *ATR1* T-DNA insertion for cross A and B and heterozygous for *ATR2* T-DNA insertion in cross C and D, indicating success in the crosses. These plants were genotyped for *CYP81A12* presence by PCR and plants that had the gene were selected for self-pollination to generate F2. F2 plants segregated 1:2:1 for *ATR1* T-DNA insertion for cross A and B and for *ATR2* T-DNA insertion for cross C and D (Tab. **S3**). Homozygous mutants were submitted for PCR amplification of *CYP81A12* and when confirmed, samples were submitted for ddPCR. Four, one, three, and five plants out of 15, 12, 9 and 15 homozygous had *CYP81A12*. These plants were analyzed by ddPCR. When only one gene copy was identified in comparison to *ALS* gene, the plants were self-pollinated, and their progenies were analyzed again using ddPCR.

The ddPCR results indicated the same copy number of *CYP81A12* and *ALS* for the parental transgenic line *CYP81A12* (Tab. **S4**). Plants selected based on previous genotyping (as indicated above from crosses A, B, C, and D) had a ratio of *ALS* and *CYP81A12* from 0.9 to 1.1, suggesting two copies of each gene, one copy per haploid genome. These plants were selected and used for further experiments.

### *atr1* and *atr2* double knockouts

Genotyping analysis of F2 plants obtained from various crosses between *atr1* and *atr2* mutants demonstrated an absence of double knockout genotypes (Tab. **S5**). A total of 416 F2 plants were genotyped, revealing seven out of the nine possible genotypes. Among the genotyped plants, 46.2% exhibited heterozygosity for both genes, 5% displayed heterozygosity for one gene and wildtype for the other, 6.2% exhibited homozygous mutation in one gene and heterozygosity in the other, while 42.6% exhibited homozygous mutation in one gene and wildtype in the other. Notably, double wildtype and double mutant genotypes were not observed. The segregation pattern observed in the F2 generation deviated from the expected Mendelian segregation for two independent genes, indicating the presence of linked genes. Further investigation revealed that the *atr1* and *atr2* genes are localized on chromosome four of Arabidopsis and are approximately two megabases (MBp) apart, providing supporting evidence that the two genes may be linked.

### *ATR1* and *ATR2* gene expression in mutant and transgenic mutant lines

Mutant *atr1-a* showed a reduced transcription of *ATR1* when the primer set used was spanning the T-DNA insertion (Fig. **4a**); however, normal expression of *ATR1* gene was observed when primer set used was downstream of the T-DNA insertion (Fig. **4b-d**), indicating that the T-DNA insertion in 5’ UTR did not decrease *ATR1* expression in *atr1-a* line. Mutant line *atr1-b*, which had T-DNA insertion in the 4^th^ intron, had significant reduction in *ATR1* transcripts. This transcript disruption was confirmed in the *ATR1* transcript when the primer set used spanned the T-DNA insertion and a section of downstream sequence (Fig. **4b-d**).

**Fig. 4.**
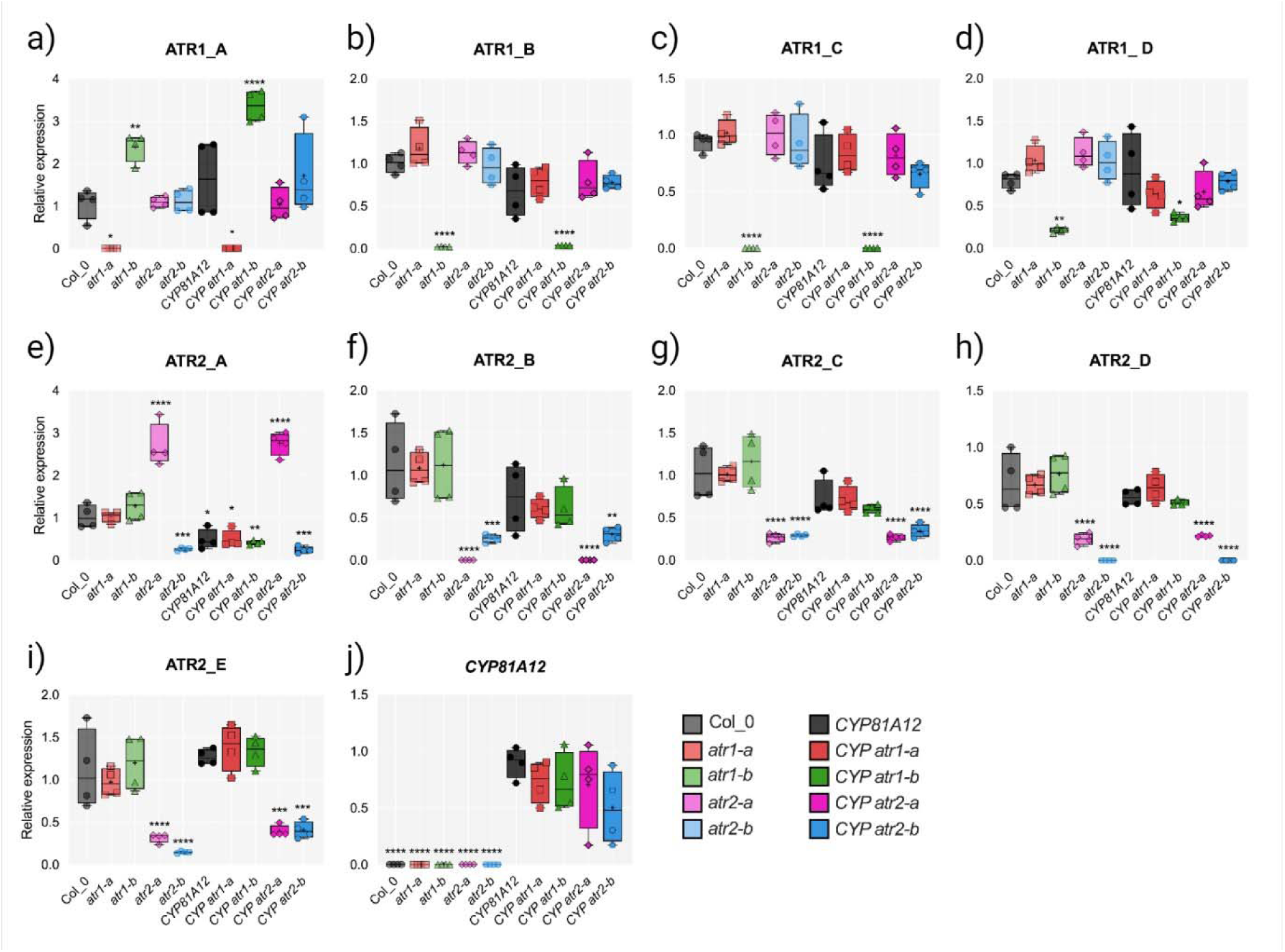
Relative gene expression of *ATR1* and *ATR2* gene in different mutant and transgenic mutant Arabidopsis lines. a-d) *ATR1* gene expression was analyzed in four different locations using different primer sets ATR1_A, ATR1_B, ATR1_C and ATR1_D. e-i) *ATR2* gene expression was analyzed in five different locations using primer set ATR2_A, ATR2_B, ATR2_C, ATR2_D and ATR2_E. j) Expression of *CYP81A12* with only one primer set. The primers set localization is illustrated in Figure 3. ATR1 and ATR2 gene expression was calculated using Col_0 as control. *CYP81A12* gene expression was calculated using transgenic Arabidopsis CYP81A12 as control. *Significant by Dunnett’s Test.

Interestingly, this line had increased transcript abundance when the primer set was upstream of the T-DNA (Fig. **4a**). These results indicate a possible compensation in the gene expression by the plant for the disrupted gene.

Mutants for *ATR2* showed similar results. Mutant *atr2-a* had an upregulation of transcript upstream of the T-DNA insertion in this line (Fig. **4e**); however, this line had significant reduction of *ATR2* expression downstream of the T-DNA insertion, especially when the set of primers used was spanning the T-DNA insertion (Fig. **4f-i**). Mutant *atr2-b* had T-DNA insertion in the 12^th^ exon and had a reduction in the transcription upstream and downstream of the T-DNA insertion (Fig. **4e-i**). T-DNA insertion in the *ATR1* mutant *atr1-b* and in the *ATR2* mutants *atr2-a* and *atr2-b* had knockdown of expression of their respective mRNA. *CYP81A12* expression was analyzed, and all the transgenic mutant lines had the same transcription production as the transgenic *CYP81A12* (Fig. **4j**).

Knocking down *ATR1* or *ATR2* did not cause obvious effects on plant growth. Transgenic *CYP81A12* had a lower fresh mass with a smaller height of the plants in comparison to Col_0 (**Fig. S2**).

### *atr1* mutant increases herbicide sensitivity in *CYP81A12* line

Based on whole-plant dose response, transgenic Arabidopsis *CYP81A12* carrying *atr1* and *atr2* were more sensitive to all herbicides tested, including penoxsulam, chlorsulfuron, mesotrione, and 2,4-D (Fig. **5**, Fig. **S4-7**). The transgenic Arabidopsis *CYP81A12* had a resistance index of 6.8, 23.6, and 20.9 for penoxsulam, mesotrione, and 2,4-D, respectively (Tab. **1**). When *ATR1* was knocked out (*CYP81A12 atr1-b*), the resistance index (RI) decreased to 1.0, 5.7, and 4.2, respectively to penoxsulam, mesotrione, and 2,4-D. The chlorsulfuron doses used in this study were not able to control the transgenic Arabidopsis *CYP81A12* and *CYP81A12 atr1-a* (Fig. **5** and Fig. **S4-7**). The resistance to chlorsulfuron of these lines was very high. The lines *CYP81A12 atr2-a* and *CYP81A12 atr2-b* showed that the highest doses imparted a slight plant control, and the model was fitted to the three-parameter model. These lines had a RI of 97.7 and 86.7 for chlorsulfuron (Tab. **1**). The line *CYP81A12 atr1-b* reduced the RI to 3.7 (Tab. **1**). *CYP81A12 atr1-b* had the highest decrease in the RI to all herbicides. *ATR2* knockdown lines (*CYP81A12 atr2-a* and *CYP81A12 atr2-b*) also reduced the RI, but not with the same magnitude as *atr1-b* (Tab. **1**).

**Fig. 5.**
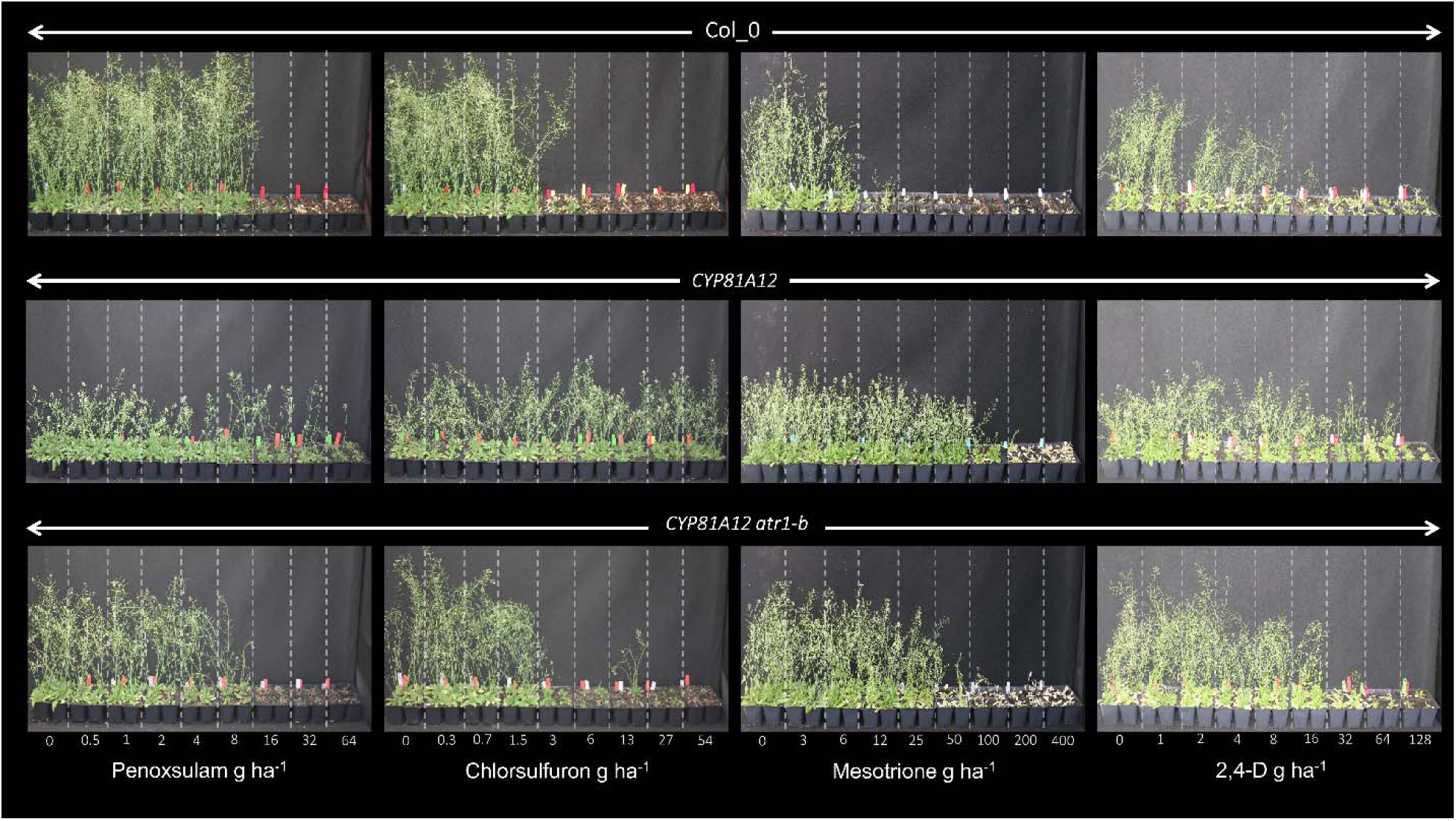
Arabidopsis lines submitted to increasing doses of penoxsulam, chlorsulfuron (ALS-inhibitors), mesotrione (HPPD-inhibitor) and 2,4-D (auxinic). Each treatment was applied on four plants, represented in the figure. Arabidopsis lines - Wild-type (Col_0), transgenic (*CYP81A12*) and transgenic mutant line *CYP81A12 atr1-b*.

**Table 1.** Parameters of the logistic equation and resistance index (RI) for shoot fresh weight (% of untreated control) of wild-type (Col_0), transgenic Arabidopsis *CYP81A12*, transgenic Arabidopsis mutants *CYP81A12 atr1-a*, *CYP81A12 atr1-b*, *CYP81A12 atr2-a* and *CYP81A12 atr2-b* treated with different doses of penoxsulam, chlorsulfuron, mesotrione and 2,4-D.

| Line | Herbicide | b | GR50 |  |  |  | RI | p-value |
| --- | --- | --- | --- | --- | --- | --- | --- | --- |
|  |  |  | Dose (g ha<sup>-1</sup>) | Lower CI (95%) | Upper CI (95%) | SE |  |  |
| Col_0 | PENO | 0.5 | 2.3 | 0.9 | 3.7 | 0.7 | - |  |
| *CYP81A12* |  | 0.7 | 15.7 | 6.7 | 24.6 | 4.5 | 6.8 | <0.001 |
| *CYP81A12 atr1-a* |  | 1.8 | 17.1 | 11.2 | 23.1 | 3.0 | 7.4 | 0.01 |
| *CYP81A12 atr1-b* |  | 0.7 | 2.3 | 1.2 | 3.5 | 0.6 | 1.0 | 0.9 |
| *CYP81A12 atr2-a* |  | 2.9 | 6.7 | 5.0 | 8.4 | 0.8 | 2.9 | 0.04 |
| *CYP81A12 atr2-b* |  | 0.7 | 3.4 | 1.5 | 5.2 | 0.9 | 1.5 | 0.4 |
| Col_0 | CHLO | 1.3 | 0.9 | 0.7 | 1.1 | 0.1 | - | - |
| *CYP81A12* |  | - | - | - | - | - | - | - |
| *CYP81A12 atr1-a* |  | - | - | - | - | - | - | - |
| *CYP81A12 atr1-b* |  | 2.4 | 3.4 | 2.5 | 4.4 | 0.4 | 3.7 | <0.01 |
| *CYP81A12 atr2-a* |  | 2.6 | 87.9 | 1.8 | 174.1 | 42.5 | 97.7 | 0.03 |
| *CYP81A12 atr2-b* |  | 0.6 | 77.5 | 1.1 | 153.9 | 37.7 | 86.7 | 0.04 |
| Col_0 | MESO | 3.7 | 4.4 | 3.85 | 5.1 | 0.3 | - | - |
| *CYP81A12* |  | 15 | 93.6 | 58.6 | 128.6 | 17.7 | 20.9 | <0.001 |
| *CYP81A12 atr1-a* |  | 5.5 | 89.3 | 78.9 | 99.7 | 5.2 | 19.9 | <0.001 |
| *CYP81A12 atr1-b* |  | 4.3 | 26.8 | 23 | 30.5 | 1.9 | 5.7 | <0.001 |
| *CYP81A12 atr2-a* |  | 2.8 | 41.8 | 35.5 | 48.1 | 3.2 | 9.3 | <0.001 |
| *CYP81A12 atr2-b* |  | 2.9 | 59.1 | 50.3 | 68.1 | 4.5 | 13.2 | <0.001 |
| Col_0 | 2,4-D | 0.5 | 5.9 | 2.7 | 9.3 | 1.6 | - | - |
| *CYP81A12* |  | 1.2 | 141.7 | 96.5 | 186.5 | 22.7 | 23.6 | <0.001 |
| *CYP81A12 atr1-a* |  | 1.6 | 103.3 | 73.7 | 132.8 | 14.9 | 17.2 | <0.001 |
| *CYP81A12 atr1-b* |  | 2.9 | 25.3 | 20.15 | 30.4 | 2.6 | 4.2 | <0.001 |
| *CYP81A12 atr2-a* |  | 0.8 | 50.1 | 29.6 | 70.5 | 10.3 | 8.3 | <0.001 |
| *CYP81A12 atr2-b* |  | 3.8 | 130.1 | 108.1 | 151.9 | 11.1 | 21.6 | <0.001 |

b: slope; d: upper limit was fixed to 100; GR50: herbicide dose that causes a 50% reduction in shoot fresh weight variable; CI: confidence interval of parameter GR50 ( $\alpha = 0.05$ ); RI: resistance index = GR50 ratio between lines with Col\_0 (control). PENO – penoxsulam, CHLO – chlorsulfuron, MESO – mesotrione.

The control of different herbicide doses on Arabidopsis lines grown in ½ MS medium yielded similar results (Fig. **6** and Tab. **2**). Among the mutants, the *CYP81A12 atr1-b* line exhibited the most significant reversal of resistance to bensulfuron, penoxsulam, chlorsulfuron, propoxycarbazone, mesotrione, and bentazon, as evidenced by the highest magnitude of response (Fig. **6** and Fig. **S7**). Notably, *CYP81A12 atr1-b* displayed symptoms similar to the Col_0 line at lower herbicide doses (Fig. **6**). In contrast, *CYP81A12 atr1-a*, *CYP81A12 atr2-a*, and *CYP81A12 atr2-b* did not exhibit reduced herbicide resistance compared to *CYP81A12*. Specifically, *CYP81A12* displayed resistance indices (RI) of 111.0, 35.7, 2.1, 4.2, 4.6, and 107.3 for bensulfuron, chlorsulfuron, penoxsulam, propoxycarbazone, mesotrione, and bentazon, respectively, while *CYP81A12 atr1-a* demonstrated reduced RIs of 22.3, 10, 0.9, 1.2, 0.9, and 0.9 for the same herbicides, reflecting similar GR_50_ values as Col_0 for penoxsulam, propoxycarbazone, mesotrione, and bentazon (Tab. **2**).

**Fig. 6.**
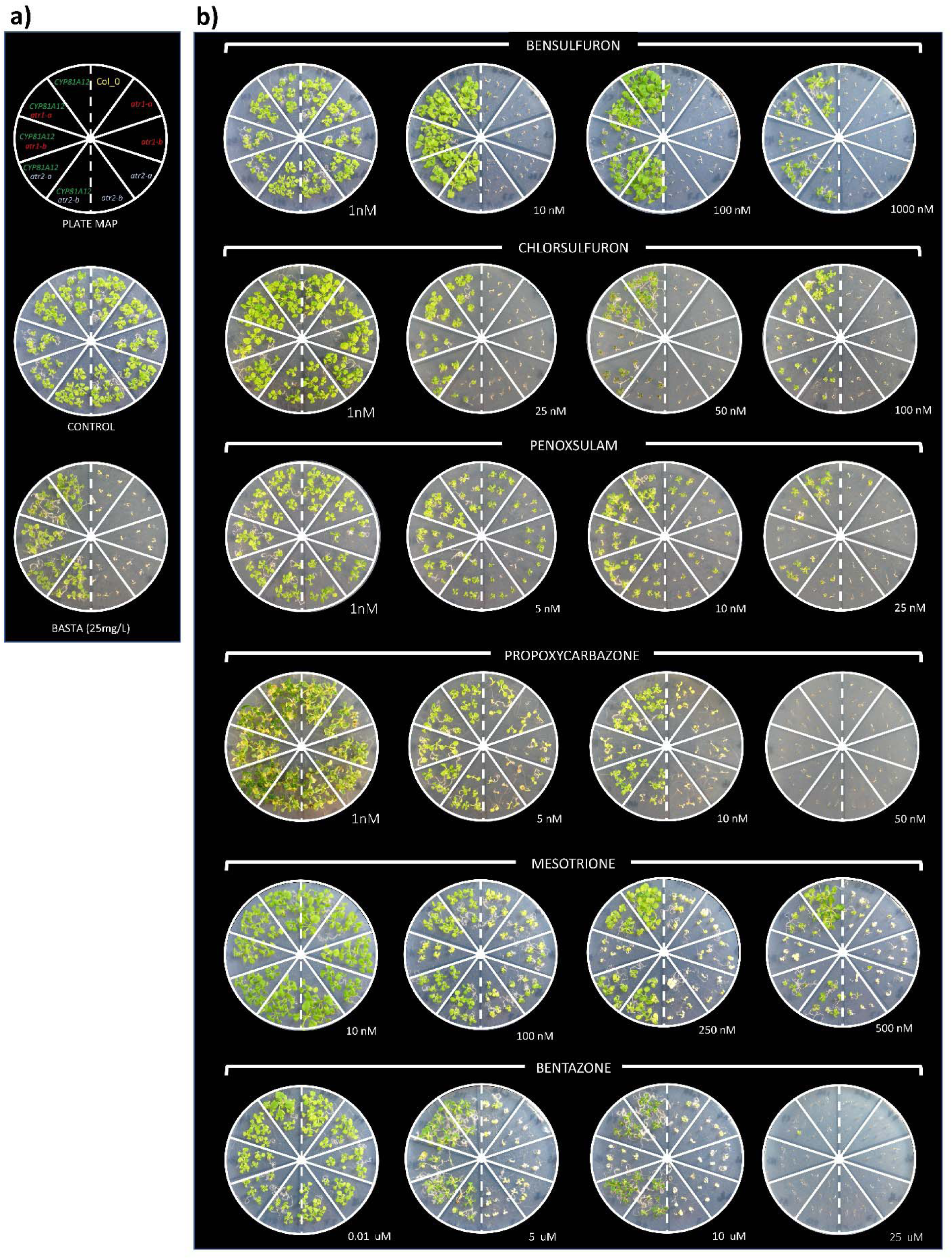
Herbicide sensitivity of wild type (Col_0), mutant *atr1-a*, *atr1-b*, *atr2-a*, *atr2-b*, transgenic Arabidopsis *CYP81A12* and transgenic mutant Arabidopsis *CYP81A12 atr1-a, CYP81A12 atr1-b*, *CYP81A12 atr2-a, CYP81A12 atr2-b*. Pictures at 14 d-old seedlings growing in ½ MS media containing different herbicide concentrations. a) Plate map, ½ MS control untreated and basta (25mg L^-1^); b) ½ MS plates with different doses of bensulfuron-methyl, chlorsulfuron, penoxsulam, propoxycarbazone, mesotrione, and bentazon.

**Table 2.**
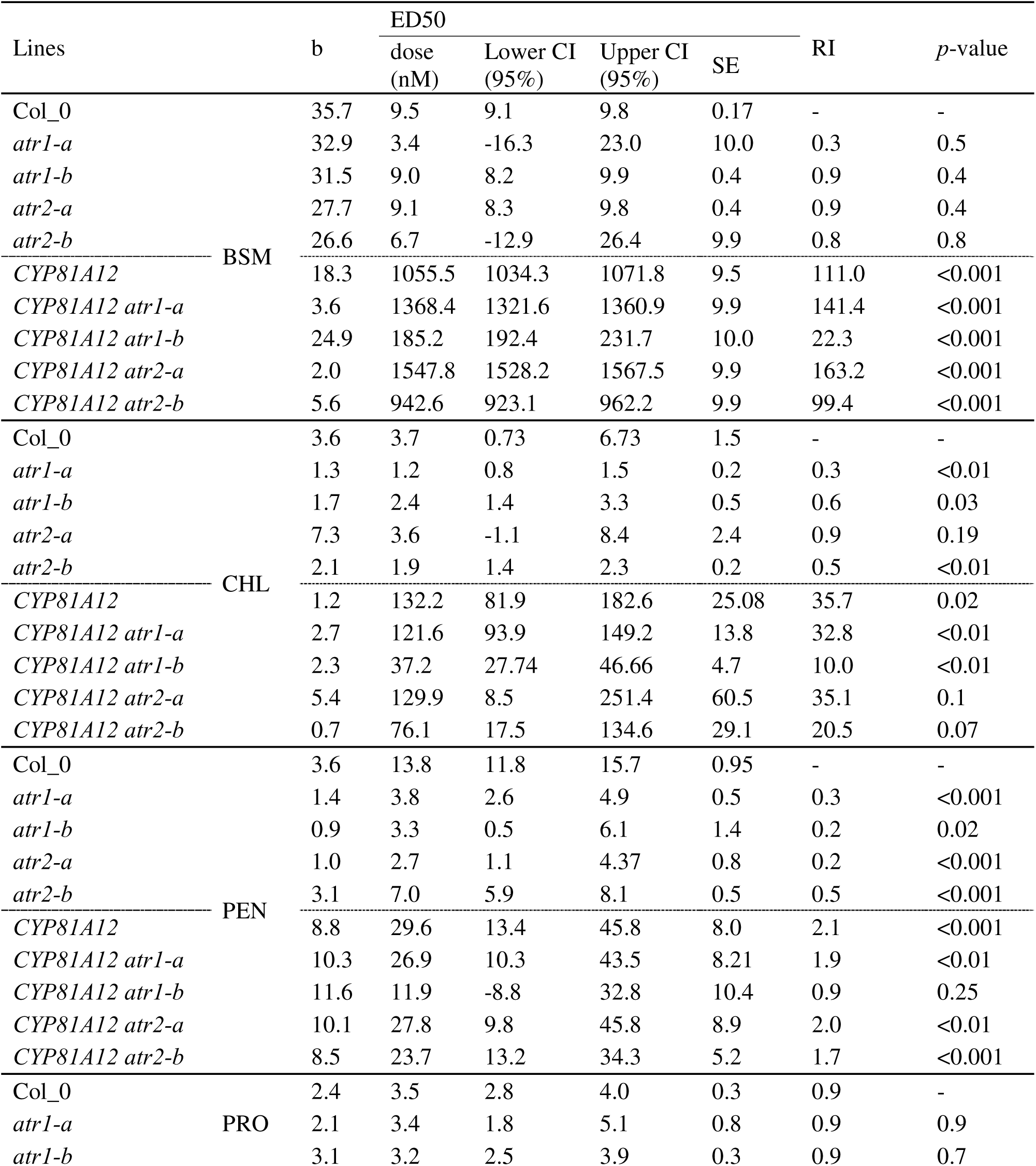
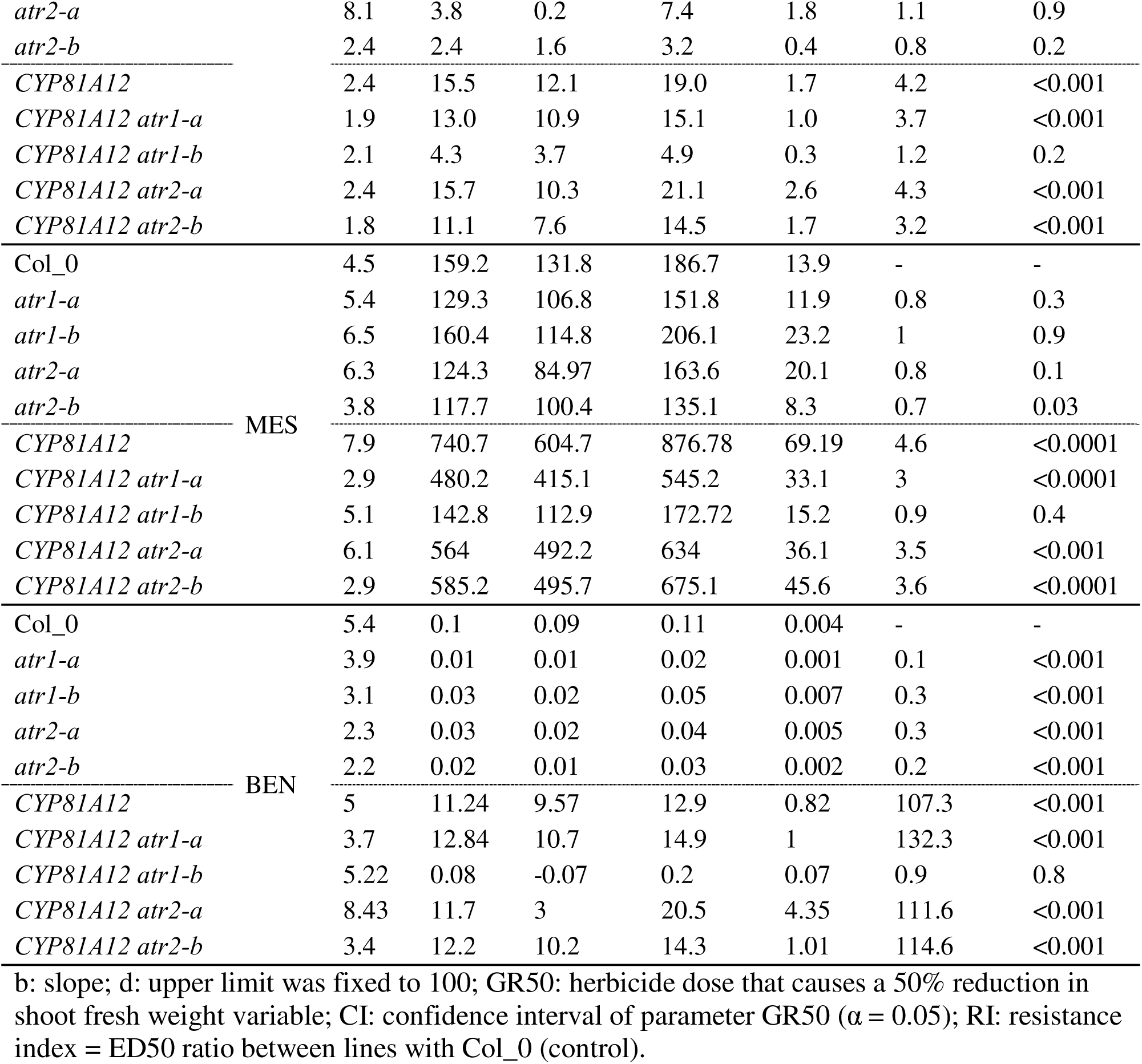
Parameters of the logistic equation and resistance index (RI) for leaf number of Arabidopsis seedlings (% of untreated control) of wild-type, *atr1-a*, *atr1-b*, *atr2-a*, *atr2-b,* and Arabidopsis transgenic *CYP81A12*, Arabidopsis transgenic mutants *CYP81A12 atr1-a*, *CYP81A12 atr1-b*, *CYP81A12 atr2-a*, *CYP81A12 atr2-b* growing in ½ MS medium supplemented with different doses of herbicides. (95%)

### Reduction of herbicide metabolism

Transgenic Arabidopsis line *CYP81A12* had a smaller peak area for the parental mesotrione than Col_0 in all time points after herbicide application, indicating faster herbicide metabolism (Fig. **7a-b**). The transgenic lines *CYP81A12 atr1-a*, *CYP81A12 atr2-a* and *CYP81A12 atr2-b* had similar profiles, with smaller peak area of parental mesotrione than Col_0. The only Arabidopsis line that showed the same peak area as Col_0 was *CYP81A12 atr1-b*.

**Fig. 7.**
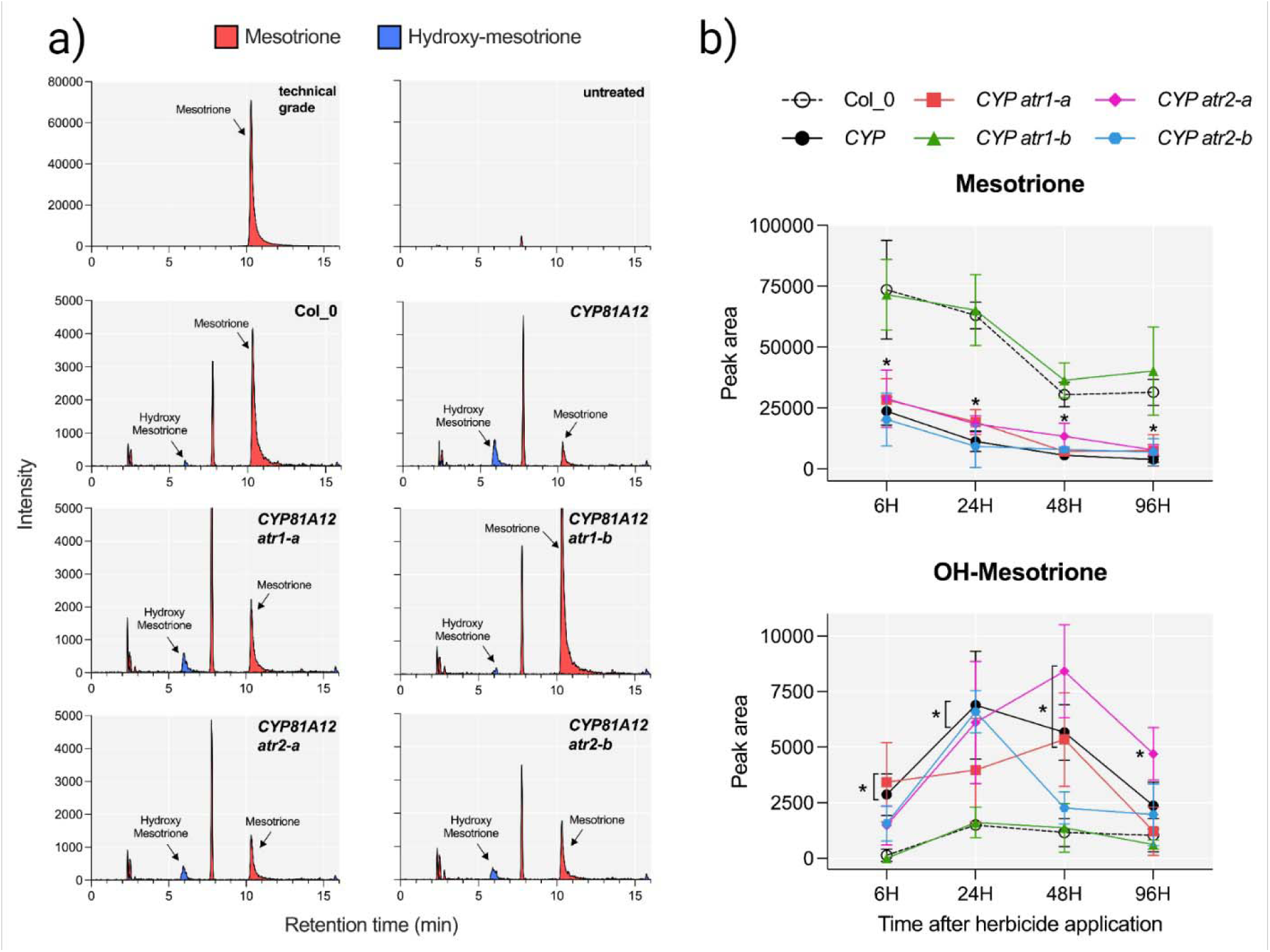
LC-MS/MS analysis of mesotrione metabolites found in different Arabidopsis lines. Arabidopsis lines analyzed were wild-type (Col_0), transgenic *CYP81A12* and transgenic mutants *CYP81A12 atr1-a*, *CYP81A12 atr1-b, CYP81A12 atr2-a, CYP81A12 atr2-b.* A) MS chromatogram 24 h after herbicide application. The retention time for parental mesotrione and hydroxy-mesotrione is 10.3- and 5.9-min, respectively. Peak at retention time 7.7 min is present in all samples, even in untreated plants. B) Peak area of mesotrione and hydroxy-mesotrione at 6, 24, 48 and 96 h after mesotrione application on different Arabidopsis lines. * *p-value* <0.05 by Dunnett’s Test.

Hydroxylation of mesotrione was higher than Col_0 for all lines in all times points, except for *CYP81A12 atr1-b* (Fig. **7a-b**), which had a very small amount of hydroxy-mesotrione, highlighting that knockdown of *ATR1* prevents mesotrione metabolism in *CYP81A12 atr1-b*. Similar results were observed for chlorsulfuron metabolism (Fig. **S8**). Transgenic *CYP81A12* had a significantly smaller peak area of parental chlorsulfuron than Col_0, inferring faster herbicide metabolism. The transgenic mutant line *CYP81A12 atr1-a* was not different in area peak of parental chlorsulfuron and hydroxy-chlorsulfuron when compared to Col_0, indicating a loss of the metabolism capacity.

### NADPH-cytochrome P450 reductase in weeds

Thirteen different weed species were analyzed, and their corresponding CPRs (cytochrome P450 reductase) were identified (Fig. **8**). The number of CPR copies varied within each weed species, ranging from two to three copies (Tab. **S6**). Through the construction of a phylogenetic tree, CPRs can be categorized into two distinct classes, namely CPR I and CPR II (Fig. **8**). These findings align with previous research, which suggests that CPR Class I is exclusive to dicot species, while Class II is present in both monocots and dicots (Jensen & Møller, 2010, Istiandari *et al*., 2021).

**Fig. 8.**
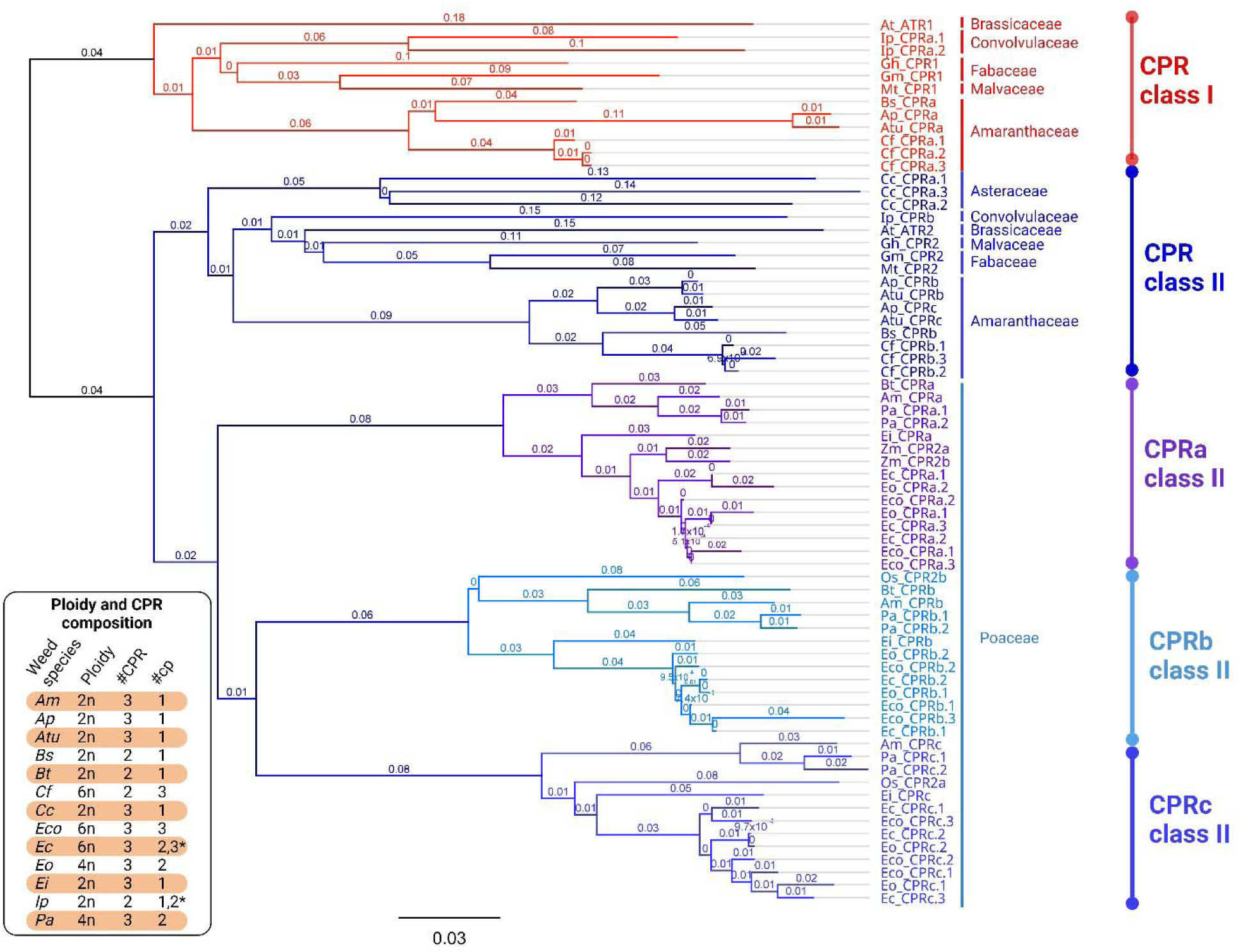
Phylogenetic tree of 69 CPR protein sequences from 13 different weed species and others 6 plant species with known CPRs sequences and classification. The phylogenetic tree shows branching of class I (red) and II (different blue shades) CPRs. Different CPR were named as “a”, “b” or “c” and the copies were numbered as “.1”, “.2” or “.3” for one, two or three copies, respectively. * Different types of CPR have different copy number in the same species. #CPR, number of cytochrome P450 reductases; #cp, number of copies. Am - *Alopecurus myosuroides* (blackgrass), At - *Arabidopsis thaliana* (Arabidopsis), Ap – *Amaranthus palmeri* (Palmer amaranth), Atu – *Amaranthus tuberculatus* (waterhemp), Bs - *Bassia scoparia* (kochia), Bt - *Bromus tectorum* (downy brome), Cc - *Conyza canadensis* (horseweed), Cf - *Chenopodium formosanum* (Djulis), Ec - *Echinochloa crus-galli* (barnyardgrass)*, Eo - E. oryzicola* (late watergrass), Eco - *E. colona* (jungle rice), Ei - *Eleusine indica* (goosegrass), Gh - *Gossypium hirsutum* (cotton), Gm - *Glycine max* (soybean), Ip - *Ipomoea purpurea* (common morning-glory), Mt - *Medicago truncatula* (Barrel medic), Os - *Oryza sativa* (rice), Pa - *Poa annua* (annual bluegrass), Zm – Zea mays (corn).

To differentiate between the CPRs within weed species, they were denoted as "a", "b", or "c" and individual copies were numbered as ".1", ".2", or ".3" to represent one, two, or three copies, respectively, in the present study. Kochia (*Bassia scoparia*), common waterhemp, common morningglory (*Ipomea purpurea*), and Djulis (*Chenopodium formosanum*) have both CPR 1 and CPR 2 (Fig. **8**). Common morning glory, a tetraploid, possesses two copies of CPR 1 and one copy of CPR 2, whereas the hexaploid Djulis has three copies of CPR 1 and three copies of CPR 2. The diploid kochia hasone copy of each CPR class. On the other hand, the diploids common waterhemp and Palmer amaranth possess one copy of CPR1 and two copies of CPR2 and horseweed possess three copies of only CPR class II. Class II Palmer amaranth CPRs have 93% identity, while the CPRs of horseweed (Class II) had a 74-76% identity among their three copies. Djulis, a hexaploid species with 27 chromosomes, exhibits three copies of each CPR class. Notably, each copy is localized within a different subgenome (B, C, and D).

Monocot plants exhibit diverse CPR compositions within CPR Class II, forming a distinct branch. Although their classification into specific classes is not documented in the literature, various monocot species possess CPR variants within Class II and we propose a division of CPR variants in grasses into CPRa, CPRb and CPRc class II (Fig. **8**). For instance, blackgrass and goosegrass each possess three different CPR II variants: Am_CPRa, Am_CPRb, and Am_CPRc for blackgrass, and Ei_CPRa, Ei_CPRb, and Ei_CPRc for goosegrass. These CPR variants are categorized into separate branches based on phylogenetic analysis. Similarly, annual bluegrass, a tetraploid species, possesses three CPR variants, with two copies of each CPR variant present in subgenomes A or B. Similarly, late watergrass, an allotetraploid species, exhibits two copies of each of the three CPR variants, distributed across its subgenomes. Jungle rice, an allohexaploid species, possesses nine CPR variants, with copies of each CPR variant located in subgenomes D, E, or F. On the other hand, barnyardgrass, also an allohexaploid species, diverges from jungle rice as it possesses eight CPR variants. Among these variants, two CPRs (CPRa and CPRc) have three copies each, while one CPR (CRPb) has two copies. It is noteworthy that the diploid downy brome is the sole monocot weed species analyzed in this study with only two CPR variants, whereas the remaining monocot weed species all possess three distinct CPR variants with different copy numbers.

All CPR genes had a well conserved domain (Fig. **S9**). Among the identified CPRs in weed genomes, three CPRs, namely Eco_CPRc.1, Ec_CPRc.3, and Eo_CPRc.1, stand out with significantly larger protein sizes compared to the others. These three CPRs have 1520, 1905, and 1948 amino acids, respectively, whereas the remaining CPRs range from 689 to 823 amino acids. Eo_CPRc.1, Ec_CPRc.3 had a bigger *N*-terminal and additionally Eco_CPRc.1 had a long *C*- terminal extensions matching phosphatidylinositol 3-kinase, which could be an assembly error.

Additionally, these three CPRs exhibit a similarity ranging from 87% to 94% among themselves. Additionally, there is close genetic relationship between subgenome D of jungle rice and subgenome C of barnyardgrass (Aoki & Yamaguchi, 2008, Wu *et al*., 2022). This suggests a common diploid ancestor, explaining the high similarity observed in CPRs among these three species.

## DISCUSSION

Cytochrome P450 reductases (CPR) are a class of enzymes that play a crucial role in the electron transfer system from NADPH to P450s (Mizutani & Ohta, 1998, Jensen & Møller, 2010). While there are numerous P450 genes, the number of CPR genes is much smaller (Backes & Kelley, 2003). In Arabidopsis, two CPR genes, class I *ATR1* and class II *ATR2*, have been identified. *ATR1* is consistently expressed, while *ATR2* is induced in response to stress (Mizutani & Ohta, 1998, Jensen & Møller, 2010). Since P450s are implicated in metabolic herbicide resistance in weeds, it is important to understand the specific role of P450 reductases in the herbicide-metabolizing capabilities of these P450s.

Through the generation of transgenic mutant Arabidopsis lines, i.e., transgenic Arabidopsis carrying *CYP81A12* from late watergrass able to metabolize a wide range of herbicides and with knockdown of either the *ATR1* or *ATR2* gene, our observations indicate that *ATR1* plays a predominant role in facilitating electron transfer from NADPH to *CYP81A12* in this particular case. Upon knocking down the *ATR1* gene in the transgenic *CYP81A12* plants (*CYP81A12 atr1-b*), reductions in herbicide resistance for 2,4-D and mesotrione were observed. Conversely, the *CYP81A12 atr1-a* line, with a T-DNA insertion in the 5’ UTR of *ATR1*, did not exhibit a reduction in herbicide resistance due to insufficient gene expression reduction (Fig. **4**). Knockouts of the *ATR2* gene (*CYP81A12 atr2a* and *atr2-b*) also led to a decrease in herbicide resistance, although not to the same extent as *ATR1*.

In Arabidopsis, *ATR1* is commonly referred to as *CPR1*, while *ATR2* is known as *CPR2* in other species. Among the identified CPRs in weeds, it was observed that only CPR class II is present in grasses. In our current study, the *ATR1* of Arabidopsis was the most crucial in providing electrons to *CYP81A12*, probably due to its best interaction efficiency of CPR:P450 complex or due its higher basal activity (TAIR, 2024). We hypothesize that different CPRs may exhibit varying interactions with various P450s, including those involved in herbicide metabolism. A recent investigation utilizing human CPRs indicated that mutations in the FMN binding domain significantly enhanced CYP activity. This suggests that amino acid variations within this domain play a significant role in the diverse interactions between CPRs and P450s (Esteves *et al*., 2020). Another study indicates that distinct CPR classes sourced from various leguminous plants (such as *Medicago truncatula*, *Lotus japonicus*, and *Glycyrrhiza uralensis*) exhibit diverse interactions with distinct P450 enzyme families within transgenic yeast.

Consequently, this variation leads to differing levels of hydroxylation activity on the substrate. (Cheng *et al*., 2023). Based on our results, it is evident that the presence of CPR in weed species exhibits variation. Notably, all examined grasses possess at least three different homologs of CPRs class II, CPRa, CPRb and CPRc, except for downy brome, which has two. One of the three different CPR class II homologs might have a higher activity and contribute the most towards reducing certain P450 genes in these species. This discrepancy in the number of CPRs among grasses suggests duplication events that occurred prior to species differentiation within the grass family (Jensen & Møller, 2010). A detailed analysis of protein sequences revealed substantial differences primarily in the *N*- and *C*-terminal regions. However, the FMN-binding and FAD-binding domains in all CPRs displayed a high degree of conservation due to their specialized functions (Fig. **S9**) and because of that, it is hypothesized that electrostatic interactions are considered to play a crucial role in guiding the formation of P450:CPR complexes.

The presence of varying numbers of CPRs in different weed species suggests distinct genetic pressures these weeds have encountered over time, leading to the acquisition of diverse functions. If herbicide application has selected resistant plants with enhanced herbicide-metabolizing activity, we hypothesize that herbicide use may have also selected for higher CPR activity to support the increased activity of P450s by facilitating electron transfer; however, this aspect needs further investigation. Additionally, in our study, we observed that in Arabidopsis, *ATR2* is unable to complement the electron transfer to *CYP81A12* when *atr1* is knocked out.

However, this may not be the case for other weed species that possess different CPR isoforms. For instance, jungle rice has three CPR isoforms, each with three copies. These CPRs could potentially be paralogs with distinct functions, interacting differentially with various P450 enzyme families, or they may have complementary functions.

Cytochrome P450 monooxygenases play pivotal roles in both primary and secondary metabolism in plants (Werck-Reichhart & Feyereisen, 2000). This significance is exemplified by studies such as the one involving homozygous mutants for CYP51A2, which exhibit defects in early embryogenesis, underscoring the enzyme’s critical role in plant growth and development (Kim *et al*., 2005). In our study, we were not able to generate *ATR1* and *ATR2* double knockouts through crosses, likely due to their inherent linkage. The hypothetical scenario of *ATR1* and *ATR2* double knockout raises intriguing possibilities, suggesting the hypothesis that a double mutation might render plants inviable due to reduced P450 activity. For most weed species examined in this study, the CPRs are dispersed across different chromosomes and do not exhibit linkage. Nevertheless, our laboratory is actively pursuing the generation of a double knockout Arabidopsis line using CRISPR/Cas 9 to gain further insights into the role of *ATR1* and *ATR2* in metabolic resistance mechanisms in weeds.

Research on the role of CPR in the evolution of pesticide resistance has primarily focused on insects, with limited investigations into the functions of CPRs in plants. In insect studies, inhibiting CPR in deltamethrin-resistant *Cimex lectularius* increased susceptibility to deltamethrin, indicating that CPR could be targeted to reduce P450 activity (Zhu *et al*., 2012). Similarly, downregulating CPR in resistant *Tetranychus cinnabarinus* (carmine spider mite) and *T. urticae* (spider mite) through RNA interference reduced P450 activity and increased susceptibility to fenpropathrin and various acaricides, respectively (Adesanya *et al*., 2020). In plants, which possess one, two, or three CPR isoforms, no research exists regarding the diverse functions of CPRs for herbicide evolution. For instance, Arabidopsis *ATR2*, which is co-expressed with lignin biosynthetic genes, showed significant alterations in lignin composition when mutated, leading to reduced activity of P450 enzymes such as cinnamate 4-hydroxylase, p-coumarate 3-hydroxylase, and ferulate 5-hydroxylase (Ehlting *et al*., 2005, Sundin *et al*., 2014). The crystal structure of Arabidopsis *ATR2* revealed that mutations in the interflavin electron transfer region can impact ATR2 activity (Niu *et al*., 2017), offering opportunities to exploit these reliable enzymes for reducing P450 activity against herbicide metabolism.

Protein alignments revealed the presence of conserved domains shared across various CPRs, along with significant variation in the FMN binding domain (Fig. **6**). There were conserved regions among CPRs in both dicot and monocot weeds, presenting a valuable opportunity to target and reduce P450 activity. Inhibiting CPR activity, either through chemical inhibitors or RNA-targeting approaches, offers a potential means to restore herbicide effectiveness against weeds exhibiting metabolic herbicide resistance. Combining CPR inhibition with herbicide treatments can serve as a strategy to control metabolic herbicide-resistant weed populations and, additionally, delay the evolution of metabolic herbicide resistance in sensitive weed populations.

The study of cytochrome P450 reductases and their interactions with cytochrome P450 enzymes is crucial for understanding metabolic herbicide resistance in weeds. Our findings highlight the importance of CPRs, particularly *ATR1* in this case of transgenic Arabidopsis, in transferring electrons to P450s involved in herbicide metabolism. The presence of different numbers of CPRs in weed species indicates the diverse genetic pressures these weeds have undergone. Our results suggest that CPR is a potential target to reduce herbicide metabolism. Identifying the most important CPRs in weeds that are closely involved in herbicide metabolism is very promising as a new tool for weed management using gene silencing approaches (Zabala-Pardo *et al*., 2022). Further research on CPRs and their functions in weeds will contribute to our understanding of herbicide resistance evolution and the development of effective weed management strategies.

## ACKNOWLEDGEMENTS

This work was supported by funding from Cotton Inc.

## AUTHOR CONTRIBUTIONS

CAGR, FED, and TAG designed the research. CAGR performed experiments. FED, TAG and SAI contributed experimental materials. CAGR, FED analyzed the data. CAGR, SAI, FED, and TAG wrote the paper. All authors contributed to editing and approving the paper.

## SUPPORTING INFORMATION

**Fig. S1.** Two-step PCR for genotyping mutant lines.

**Fig. S2.** Phenotype analysis of transgenic mutant lines.

**Fig. S3.** T-DNA insertion genotyping for *ATR1* (a and b) and *ATR2* gene (c and d) in different Arabidopsis mutant lines.

**Fig. S4.** Pictures of different lines of Arabidopsis submitted to different doses of mesotrione and 2,4-D.

**Fig. S5.** Pictures of different lines of Arabidopsis submitted to different doses of penoxsulam and chlorsulfuron.

**Fig. S6.** Different lines of Arabidopsis in response to increasing doses of different herbicides.

**Fig. S7.** Dose-response of different lines of Arabidopsis growing in different herbicides concentration in ½ MS plates.

**Fig. S8.** LC-MS/MS analysis of chlorsulfuron metabolites found in different Arabidopsis lines.

**Fig. S9.** Multiple protein alignment of 67 CPR protein sequences from 13 different weed species and others six plant species with colors indicating conserved residue between sequences.

**Table S1.** Primers and their characteristics used to genotype by PCR *ATR1* and *ATR2* genes with the T-DNA insertion and *CYP81A12* in F1 or F2 plants.

**Table S2.** Primers and their characteristics used for gene expression analysis of *ATR1*, *ATR2*, and CYP81A12.

**Table S3.** PCR genotyping for T-DNA insertion in F2 segregating plants derived from crosses between *atr1* or *atr2* mutants with transgenic Arabidopsis *CYP81A12*.

**Table S4.** ddPCR results of parental transgenic Arabidopsis *CYP81A12* and four selected F3 plants from crosses A (*CYP81A12* X *atr1-a*), B (*CYP81A12* X *atr1-b*), C (*CYP81A12* X *atr2-a*), and D (*CYP81A12* X *atr2-b*).

**Table S5.** Genotype analysis of F2 segregating populations derived from crosses between *atr1* and *atr2* Arabidopsis mutants.

**Table S6.** Weed species and their information about NADPH-cytochrome P450 reductase composition.

