## Supporting Information for "Unraveling the Role of P450 Reductase in Herbicide Metabolic Resistance Mechanism"

The following Supporting Information is available for this article:

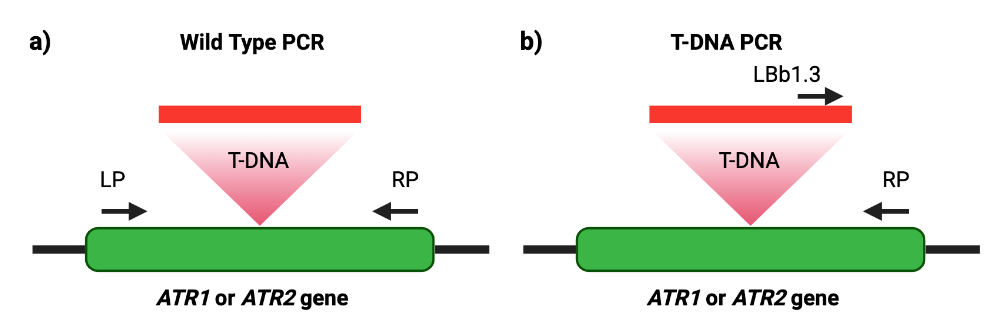

**Fig. S1.** Two-step PCR for genotyping mutant lines. The “wild-type PCR” reaction (a) amplifies a region that is present in wild type and heterozygous lines, but it does not amplify in homozygous lines. The “T-DNA PCR” (b) amplifies only if T-DNA insertion is present. LP – left primer, RP – right primer, LBb1.3 – left border primer for T-DNA insertion.

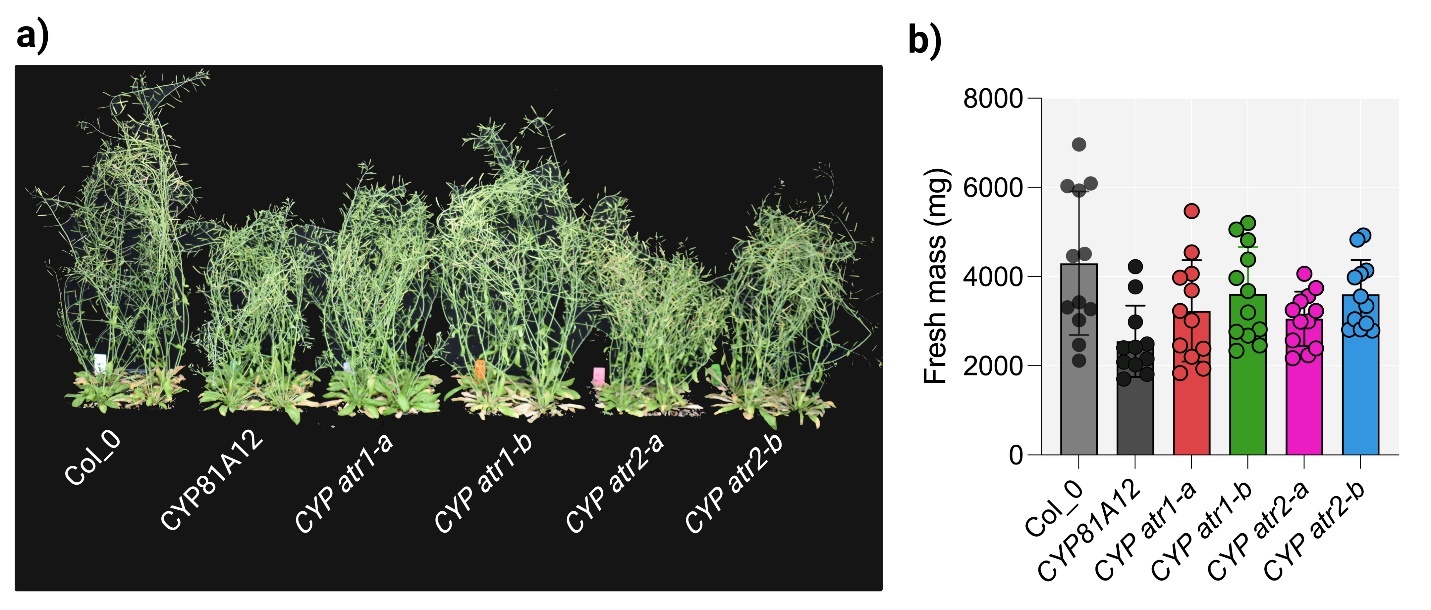

**Fig. S2.** Phenotype analysis of transgenic mutant lines. a) Picture at 28 days after transplanting of four replicates for each line. b) Fresh mass of twelve plants of each line at 28 days after transplanting.

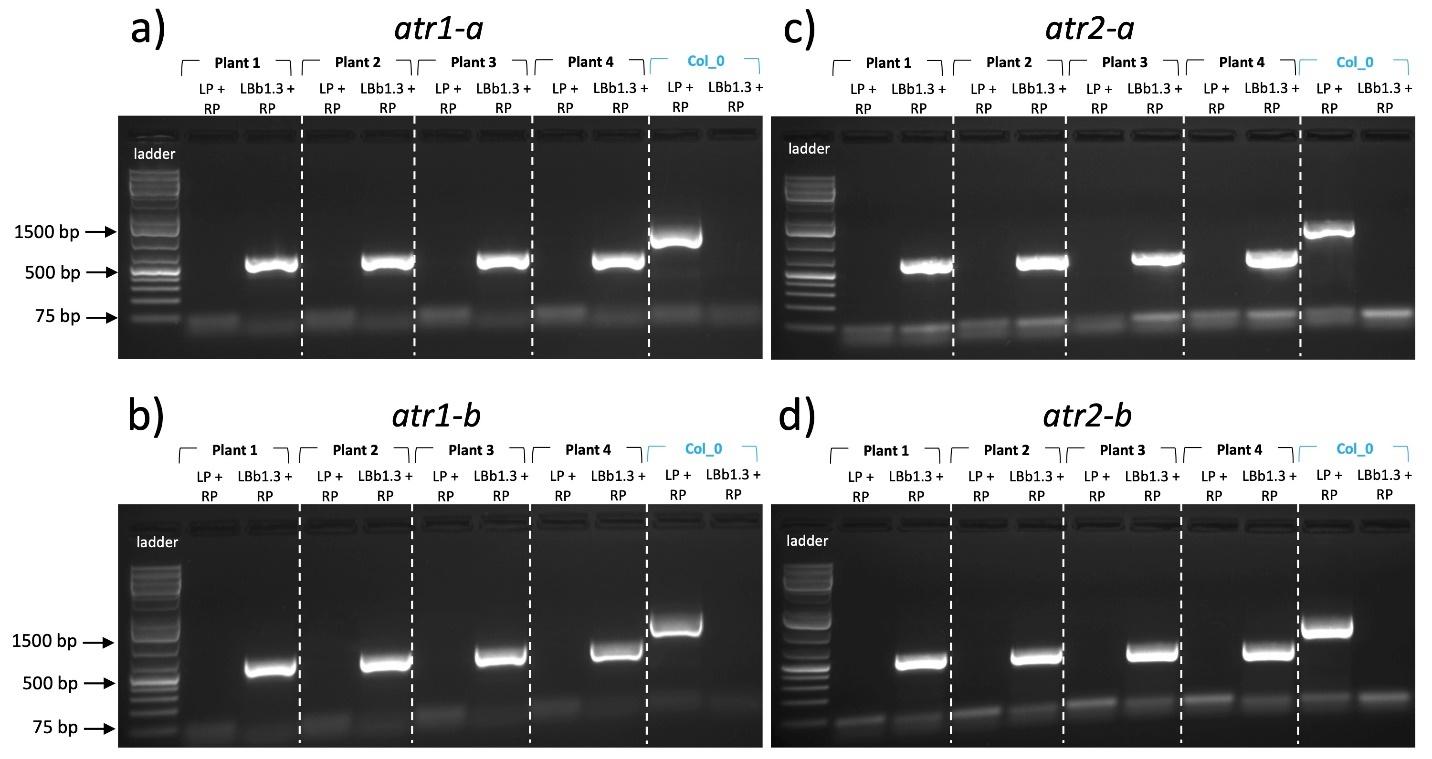

**Fig. S3.** T-DNA insertion genotyping for *ATR1* (a and b) and *ATR2* gene (c and d) in different Arabidopsis mutant lines. Each subfigure includes four sample plants of each line and one wildtype columbia_0 (Col_0) as a control. Each plant was submitted for two PCR, first reaction was “wildtype PCR” using primer set LP + RP, and second reaction “T-DNA PCR” reaction using primers set LBb1.3 + RP.

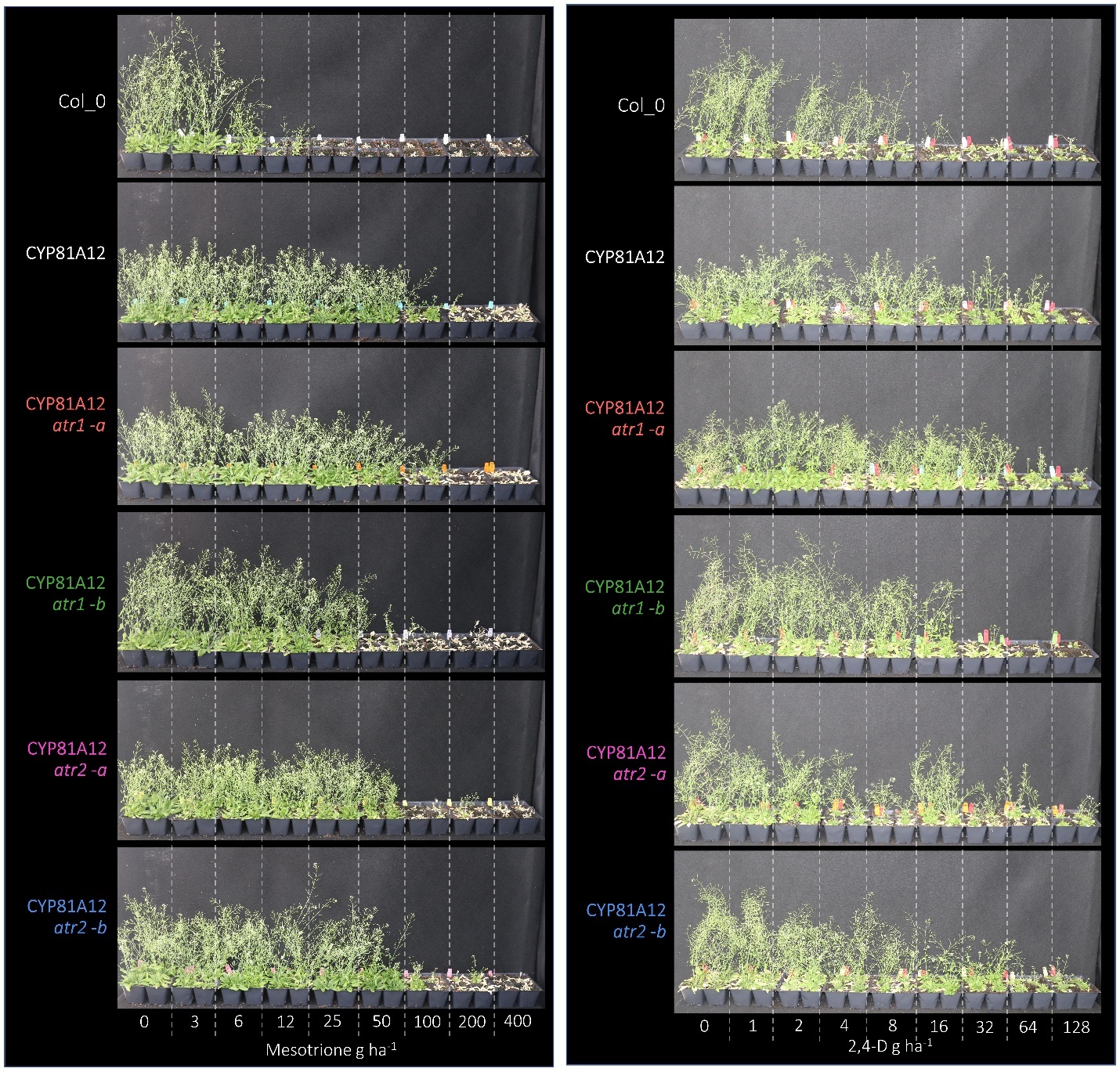

**Fig. S4.** Pictures of different lines of Arabidopsis submitted to different doses of mesotrione and 2,4-D. Pictures of four replicates at 28 d after herbicide application. Col_0, wild-type; *CYP81A12*, transgenic line expression *CYP81A12*; *CYP81A12 atr1-a*, transgenic Arabidopsis expressing *CYP81A12* carrying T-DNA insertion on *ATR1* 5’UTR; *CYP81A12 atr1-b*, transgenic Arabidopsis expressing *CYP81A12* carrying T-DNA insertion on *ATR1* 4^th^ intron; *CYP81A12 atr2-a*, transgenic Arabidopsis expressing *CYP81A12* carrying T-DNA insertion on *ATR2* 3^th^ intron; *CYP81A12 atr2-b*, transgenic Arabidopsis expressing *CYP81A12* carrying T-DNA insertion on *ATR2* 12^th^ exon.

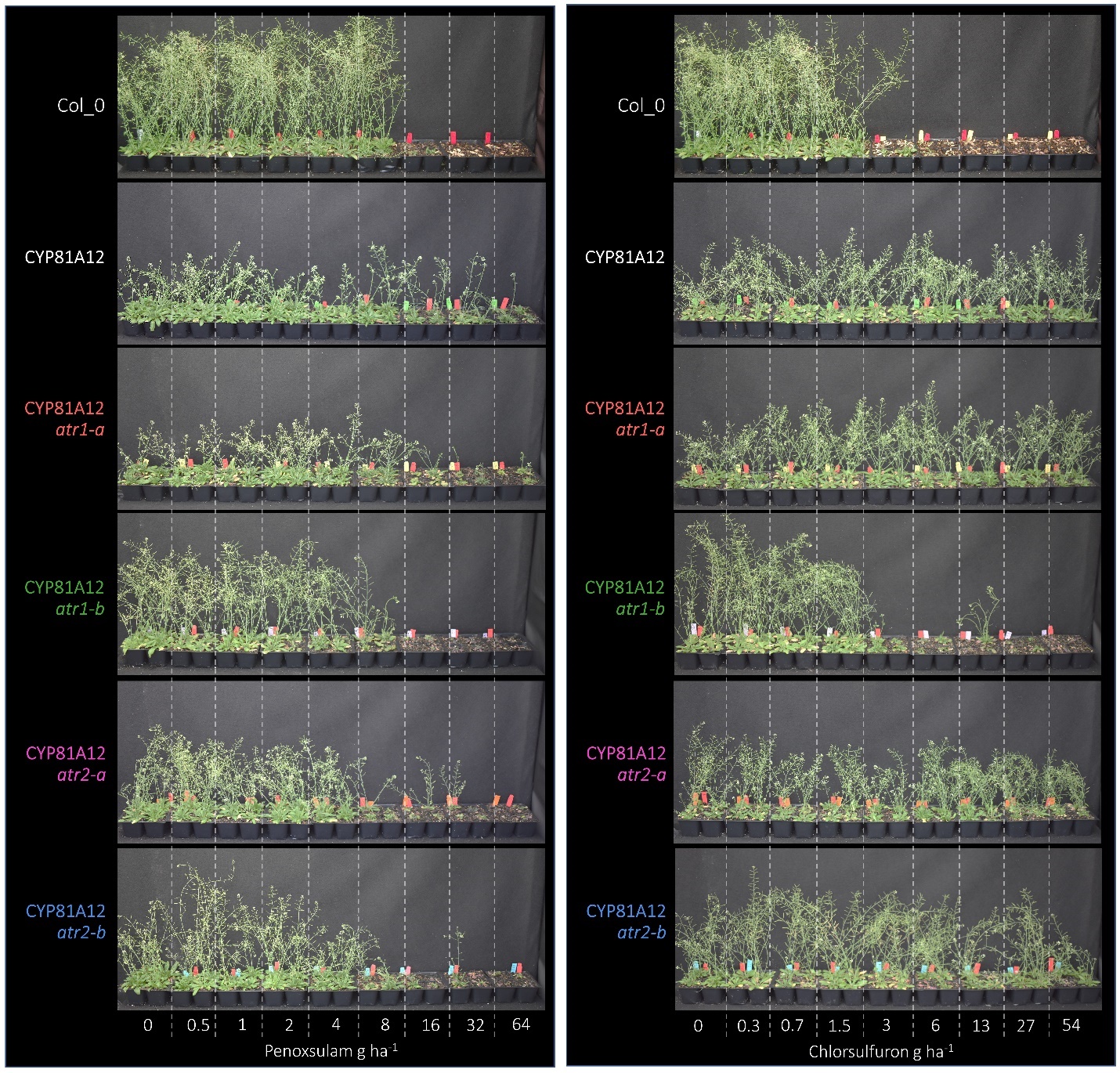

**Fig. S5.** Pictures of different lines of Arabidopsis submitted to different doses of penoxsulam and chlorsulfuron. Pictures of four replicates at 28 d after herbicide application. Col_0, wild-type; *CYP81A12*, transgenic line expression *CYP81A12*; *CYP81A12 atr1-a*, transgenic Arabidopsis expressing *CYP81A12* carrying T-DNA insertion on *ATR1* 5’UTR; *CYP81A12 atr1-b*, transgenic Arabidopsis expressing *CYP81A12* carrying T-DNA insertion on *ATR1* 4^th^ intron; *CYP81A12 atr2-a*, transgenic Arabidopsis expressing *CYP81A12* carrying T-DNA insertion on *ATR2* 3^th^ intron; *CYP81A12 atr2-b*, transgenic Arabidopsis expressing *CYP81A12* carrying T-DNA insertion on *ATR2* 12^th^ exon.

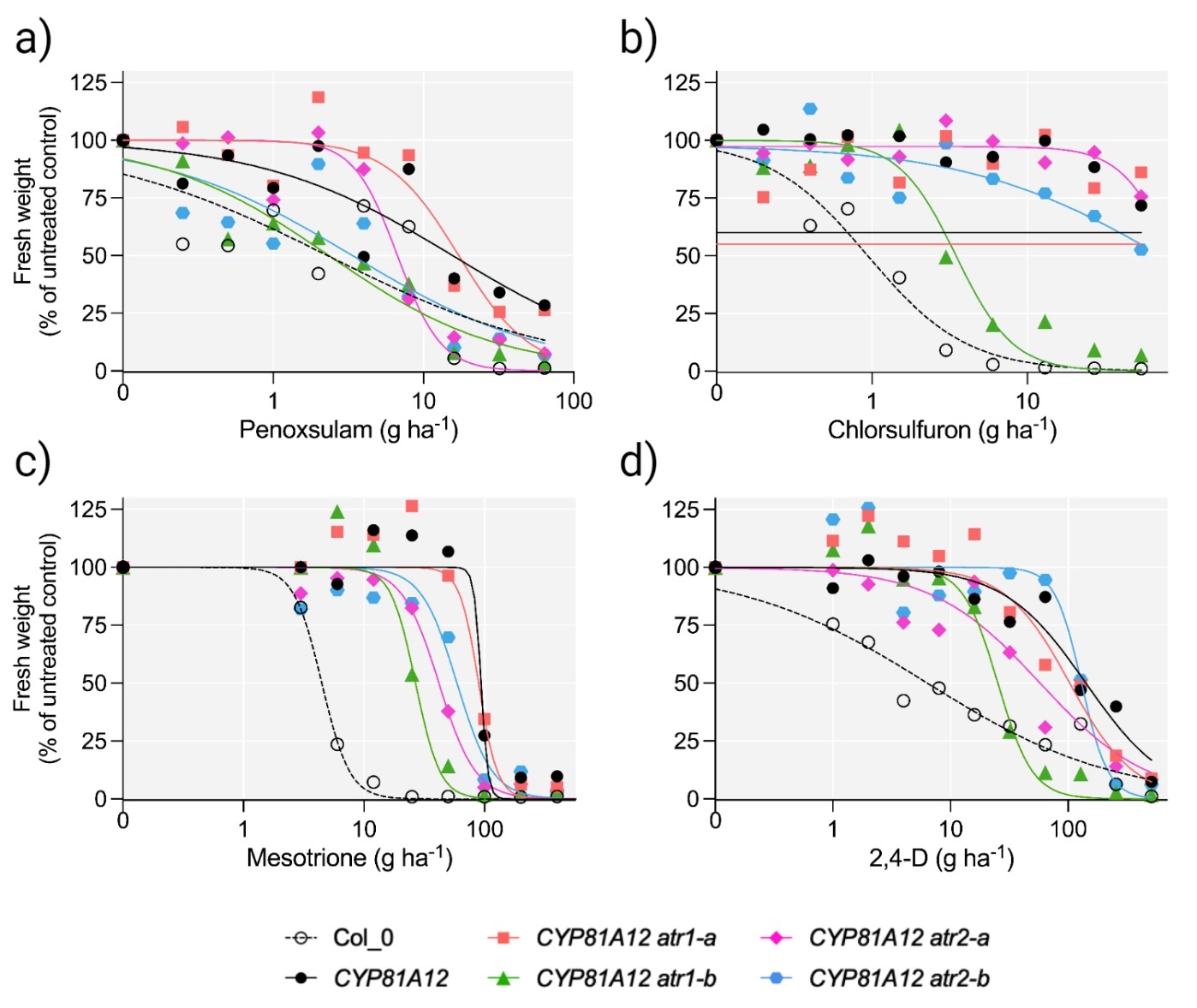

**Fig. S6.** Different lines of Arabidopsis in response to increasing doses of different herbicides. a) penoxsulam, b) chlorsulfuron, c) mesotrione and d) 2,4-D. Col_0, wild-type; *CYP81A12*, transgenic line expression *CYP81A12*; *CYP81A12 atr1-a*, transgenic Arabidopsis expressing *CYP81A12* carrying T-DNA insertion on *ATR1* 5’UTR; *CYP81A12 atr1-b*, transgenic Arabidopsis expressing *CYP81A12* carrying T-DNA insertion on *ATR1* 4^th^ intron; *CYP81A12 atr2-a*, transgenic Arabidopsis expressing *CYP81A12* carrying T-DNA insertion on *ATR2* 3^th^ intron; *CYP81A12 atr2-b*, transgenic Arabidopsis expressing *CYP81A12* carrying T-DNA insertion on *ATR2* 12^th^ exon.

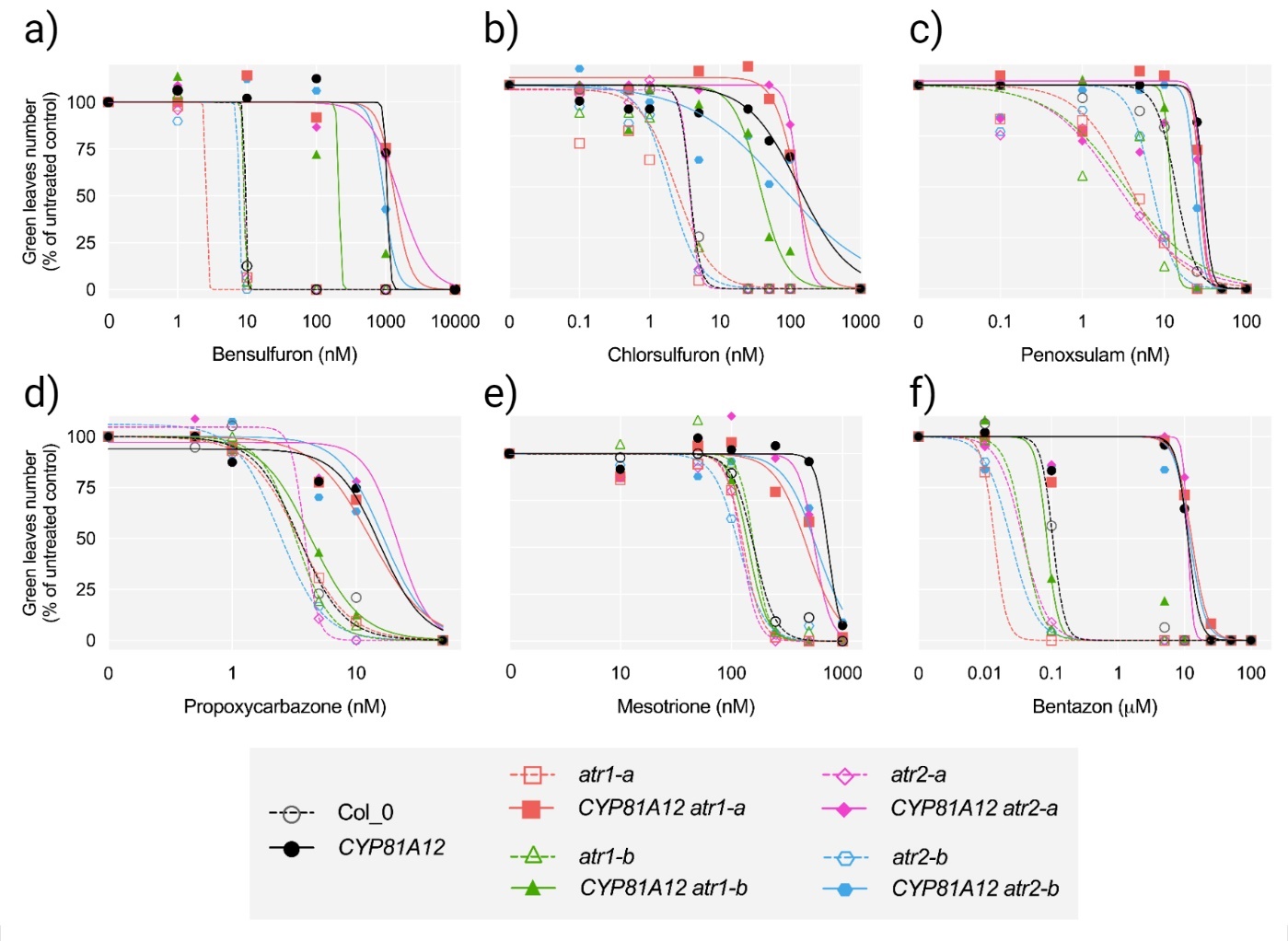

**Fig. S7.** Dose-response of different lines of Arabidopsis growing in different herbicides concentration in ½ MS plates. Arabidopsis wild-type (Col_0), transgenic *CYP81A12*, mutant lines *atr1-a*, *atr1-b*, *atr2-a* and *atr2-b* and transgenic mutant lines expressing *CYP81A12, CYP81A12 atr1-a*, *CYP81A12 atr1-b*, *CYP81A12 atr2-a* and *CYP81A12 atr2-b*. A) dose-response curve to bensulfuron, B) chlorsulfuron, C) penoxsulam, D) propoxycarbazone, E) mesotrione and F) bentazon.

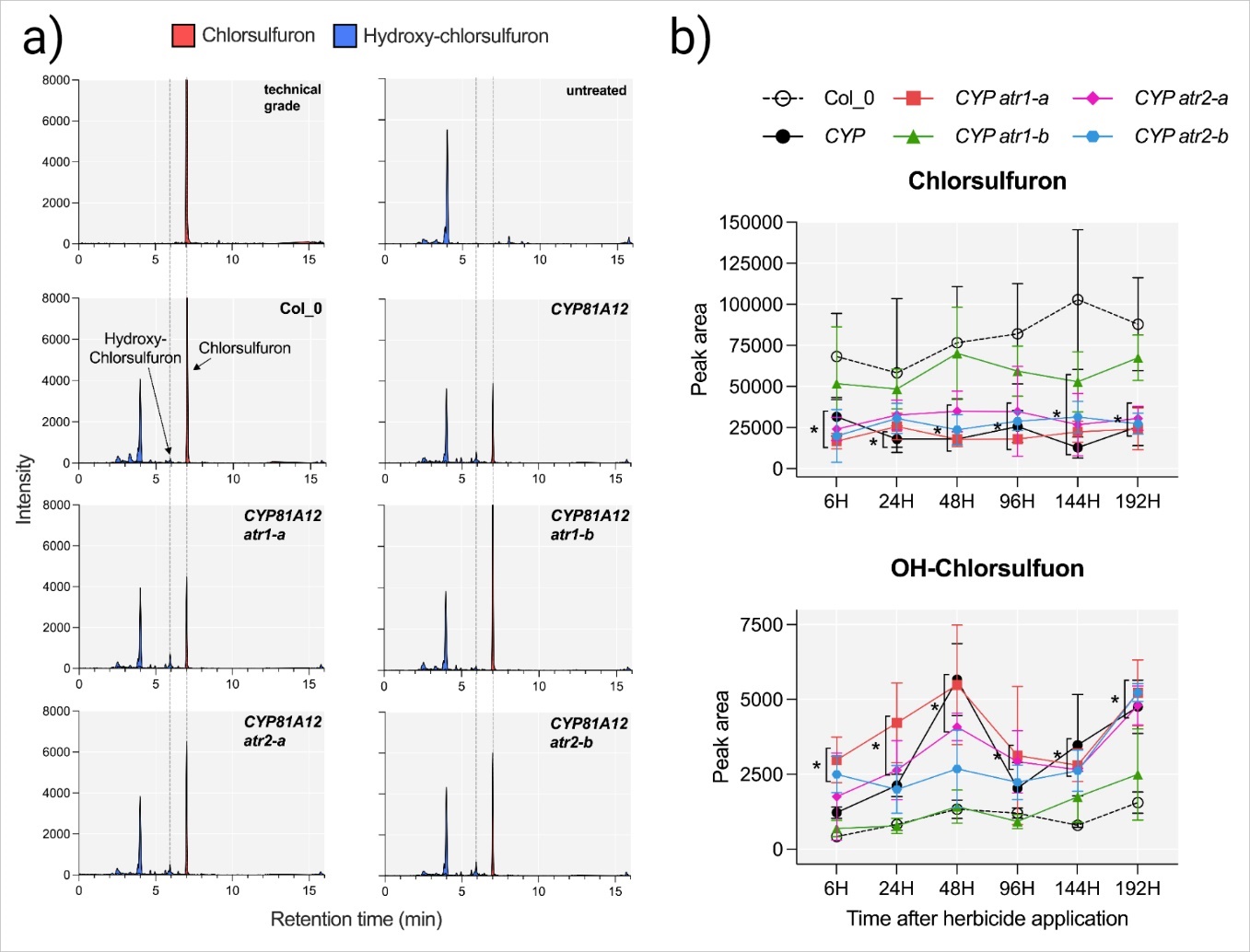

**Fig. S8.** LC-MS/MS analysis of chlorsulfuron metabolites found in different Arabidopsis lines. Arabidopsis lines analyzed were wild-type (Col_0), transgenic *CYP81A12* and transgenic mutants *CYP81A12 atr1-a*, *CYP81A12 atr1-b, CYP81A12 atr2-a, CYP81A12 atr2-b.* A) MS chromatogram 192 h after herbicide application. The retention time for parental chlorsulfuron and hydroxy-chlorsulfuron 7.01 and 5.9 min, respectively. Peak at retention time 4.01 min is present in all samples, even in untreated plants. B) Peak area of chlorsulfuron and hydroxy-chlorsulfuron at different time points after chlorsulfuron application. * *p-value* <0.05 by Dunnett’s Test.

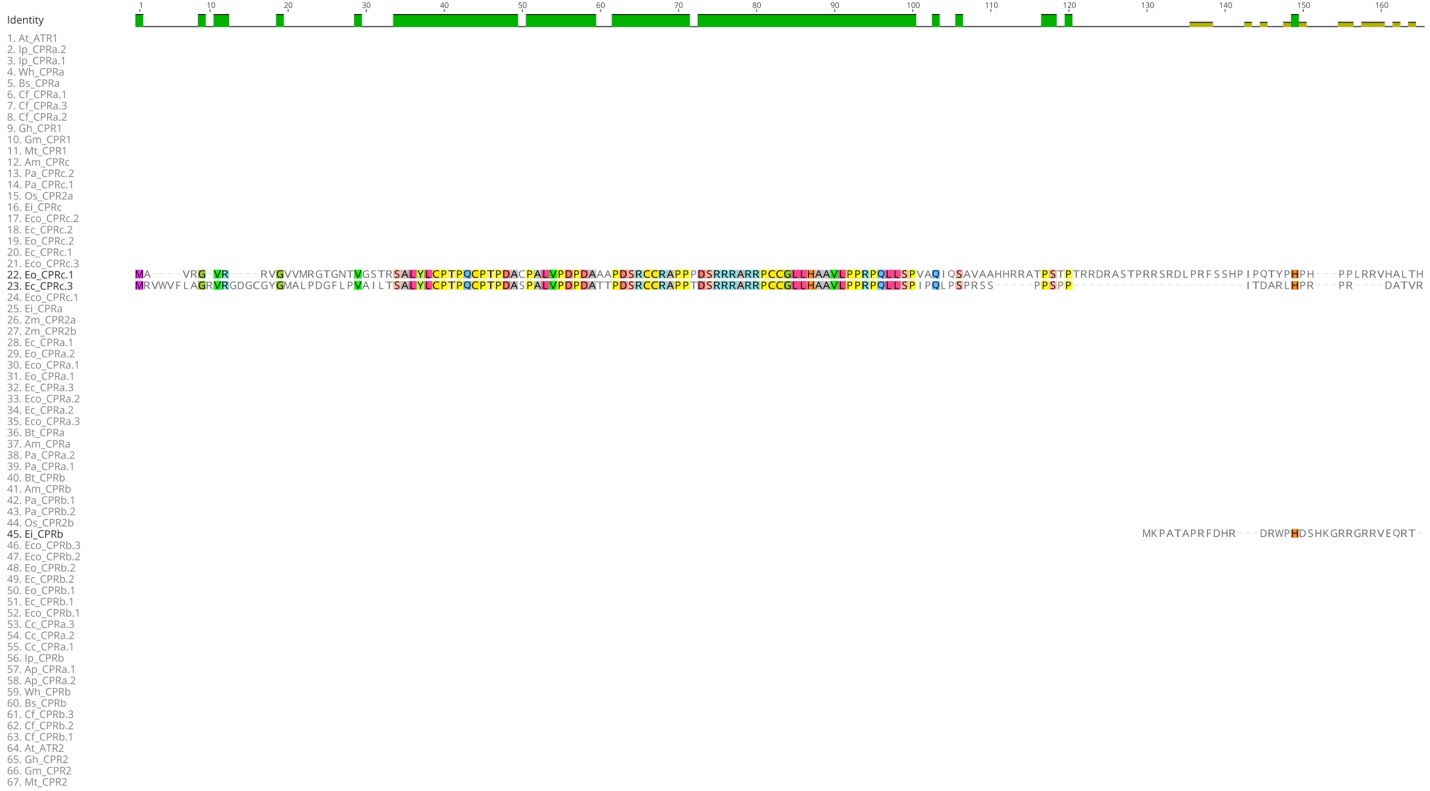

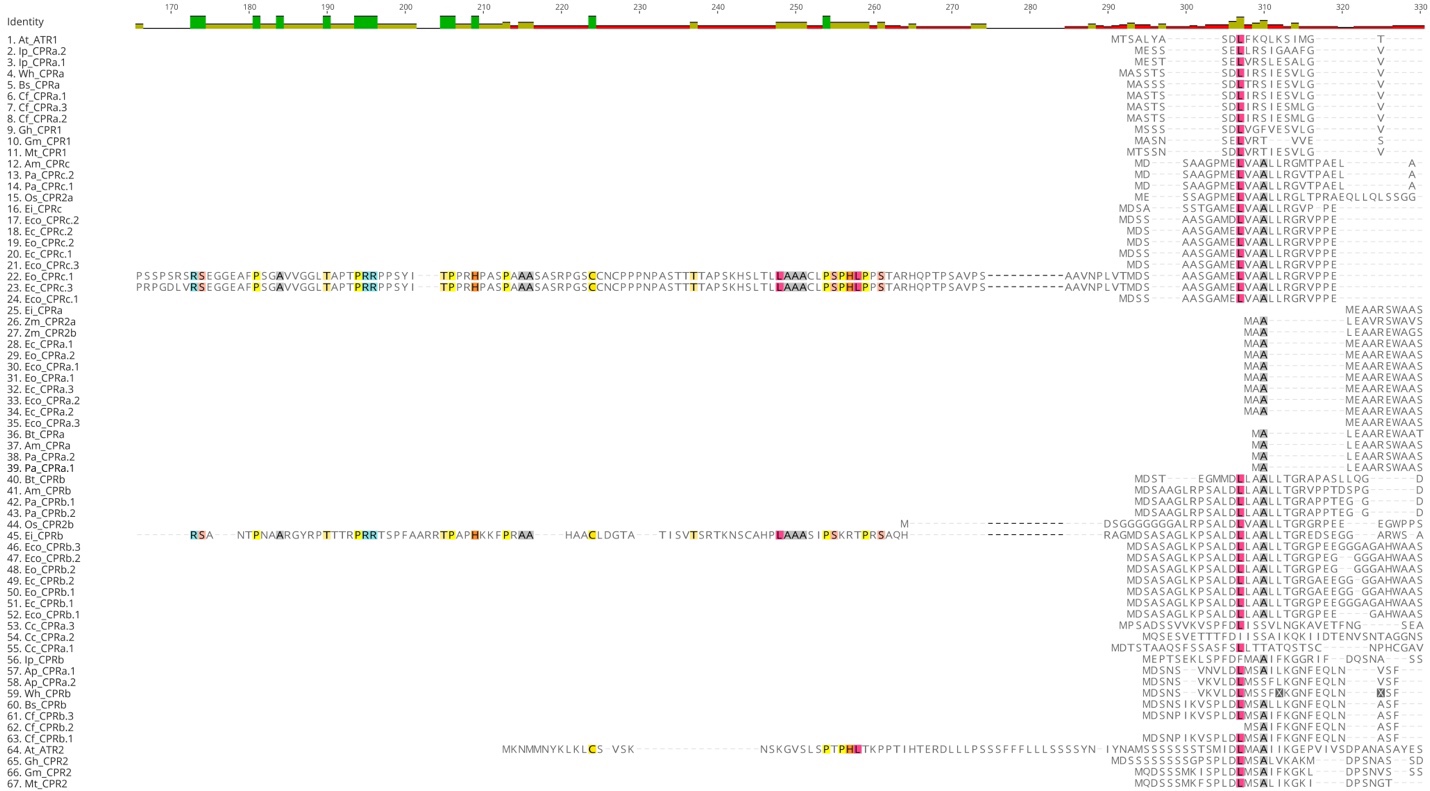

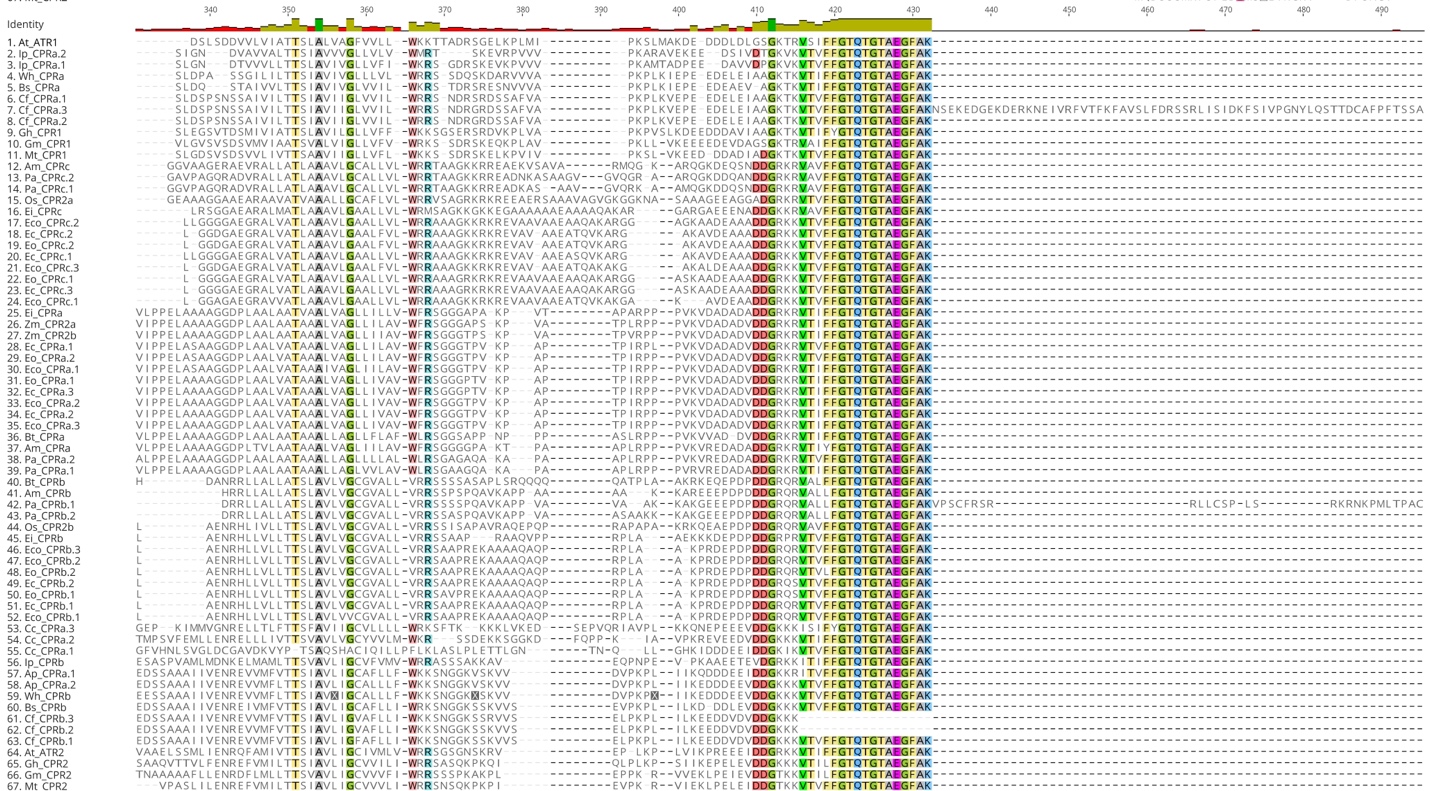

FMN binding

Transmembrane

FMN binding

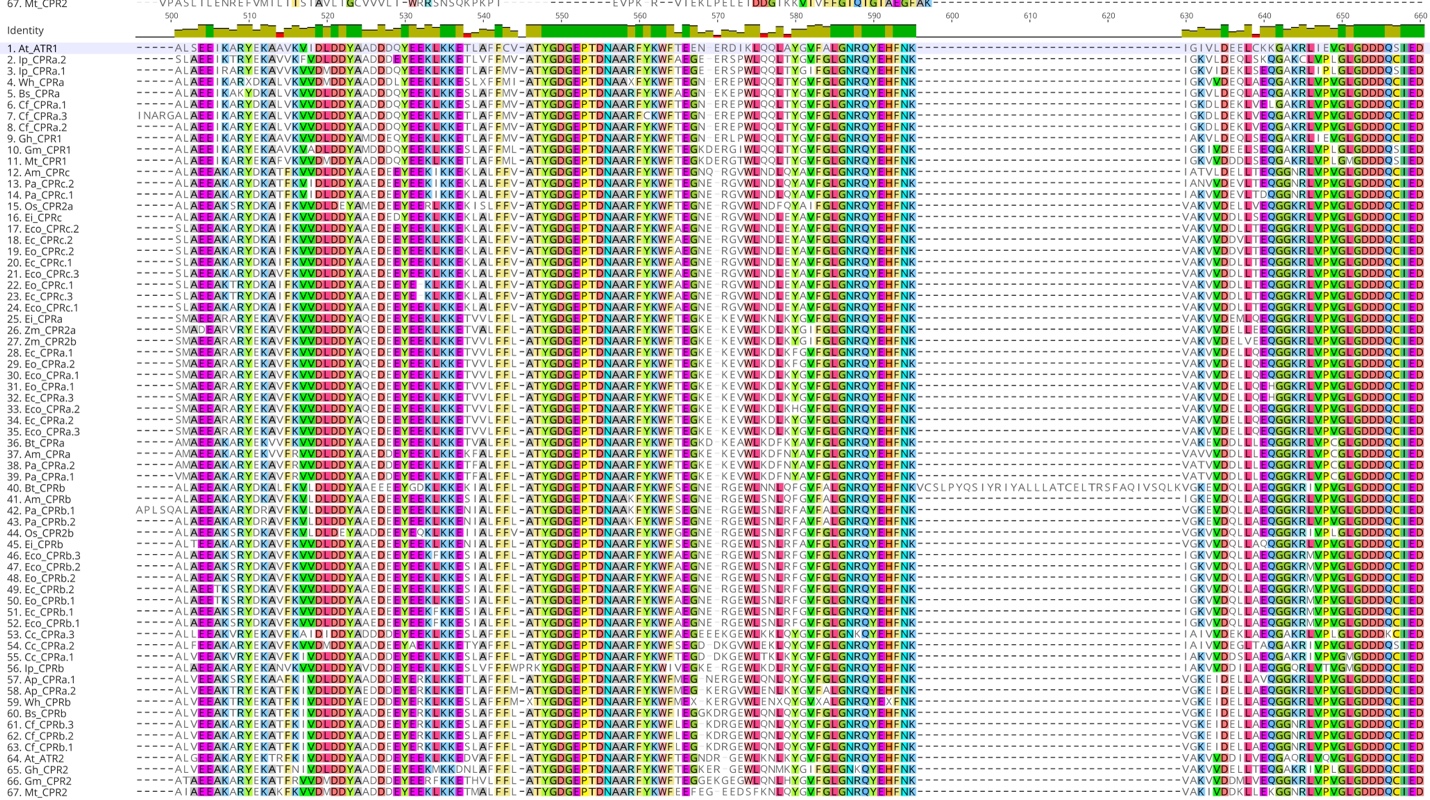

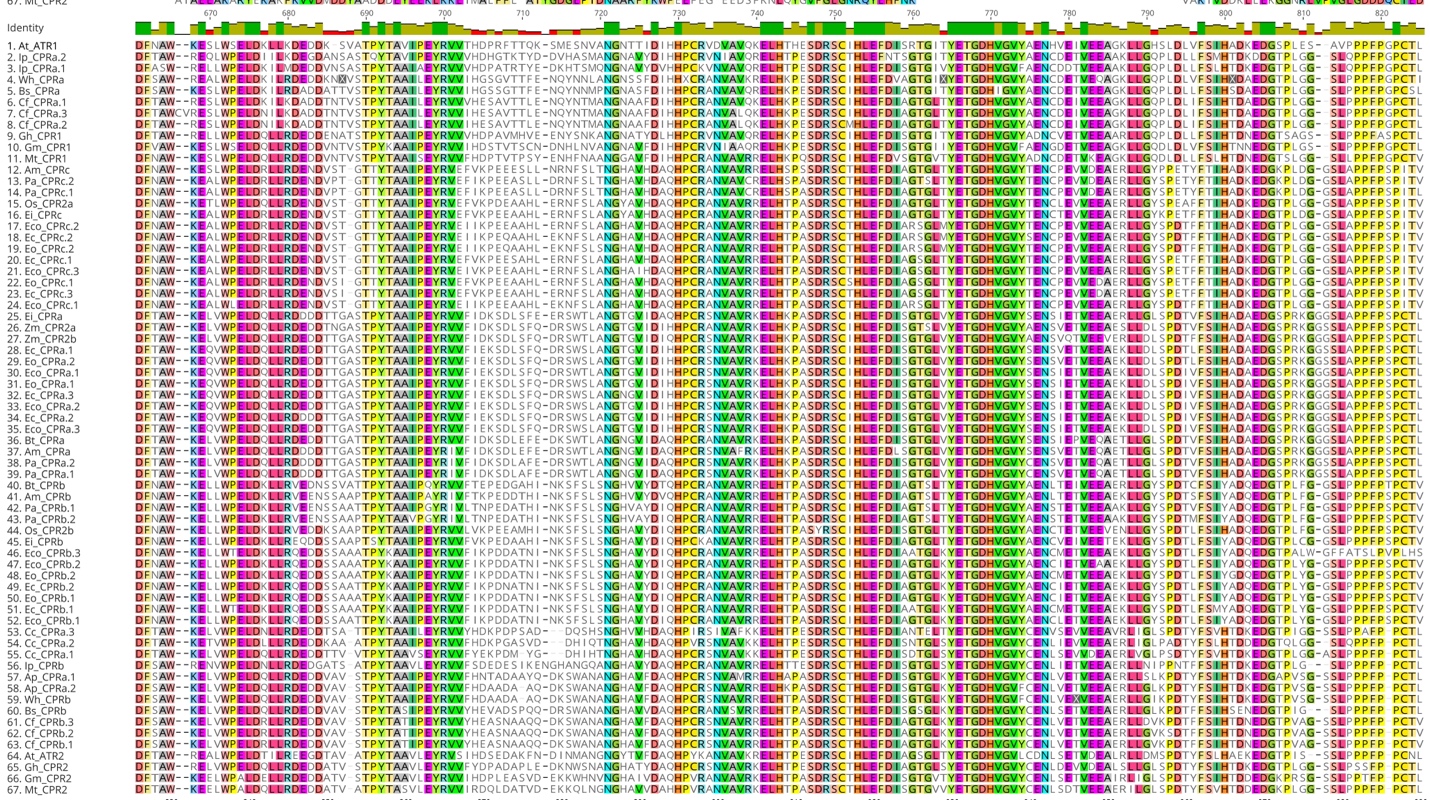

FAD binding

FAD binding

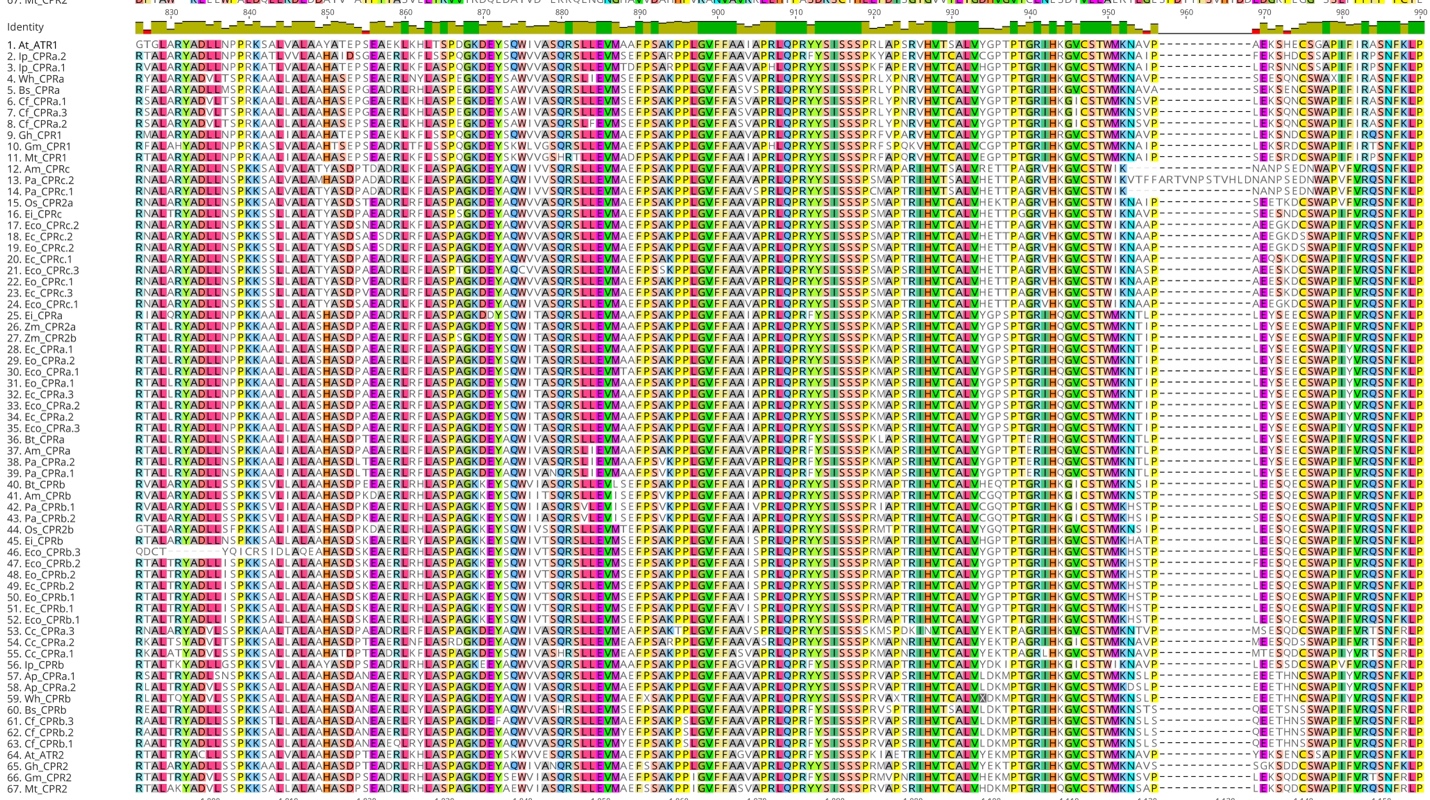

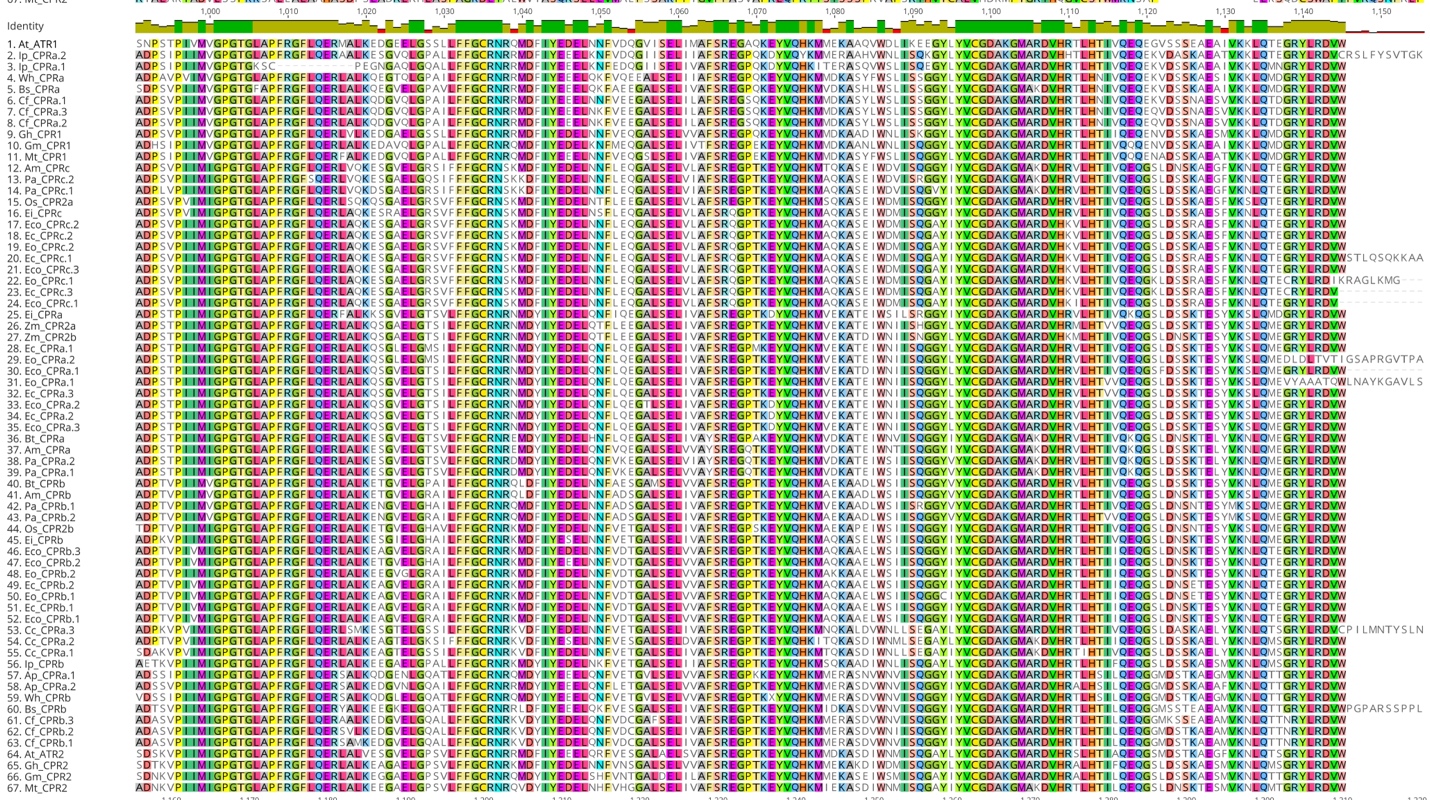

NADP binding

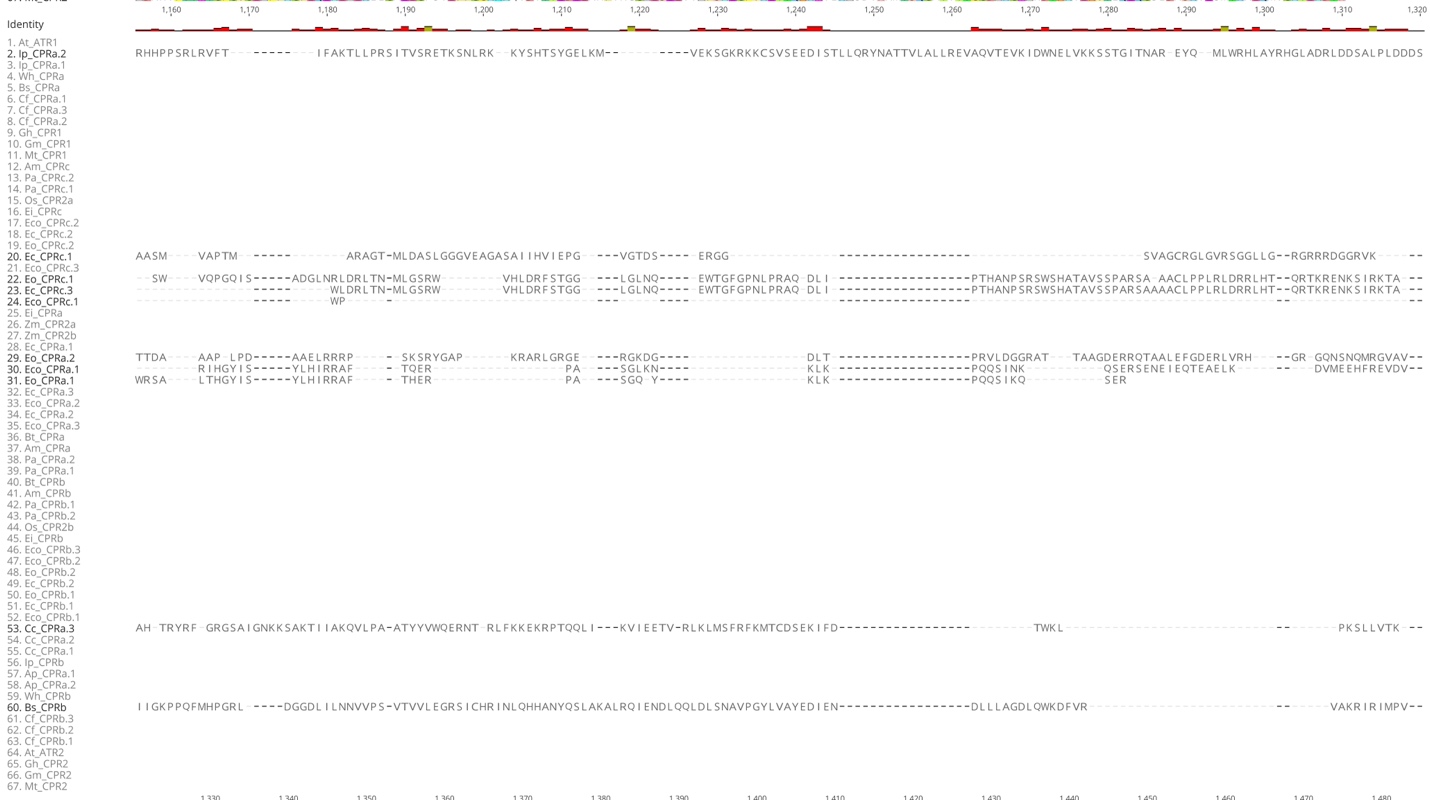

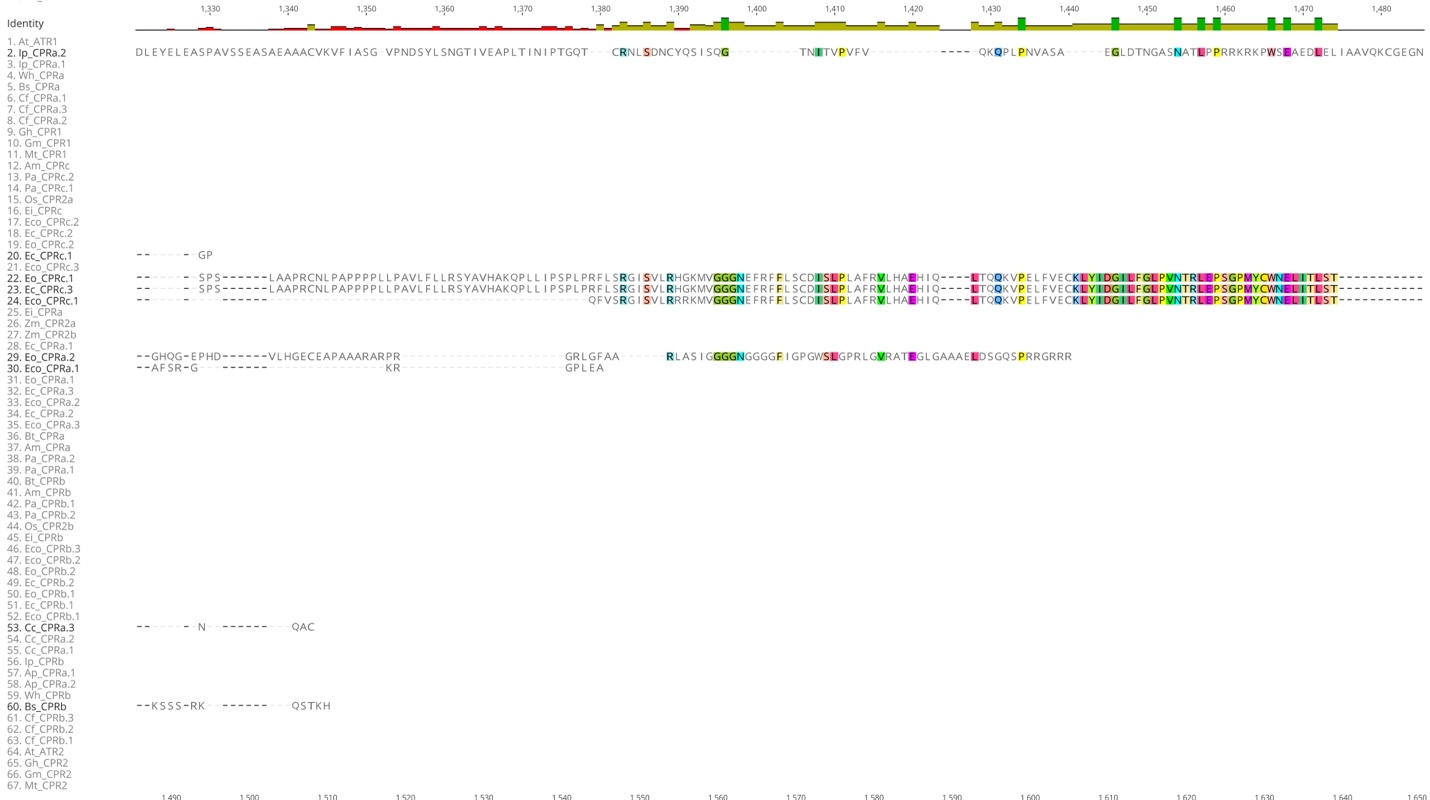

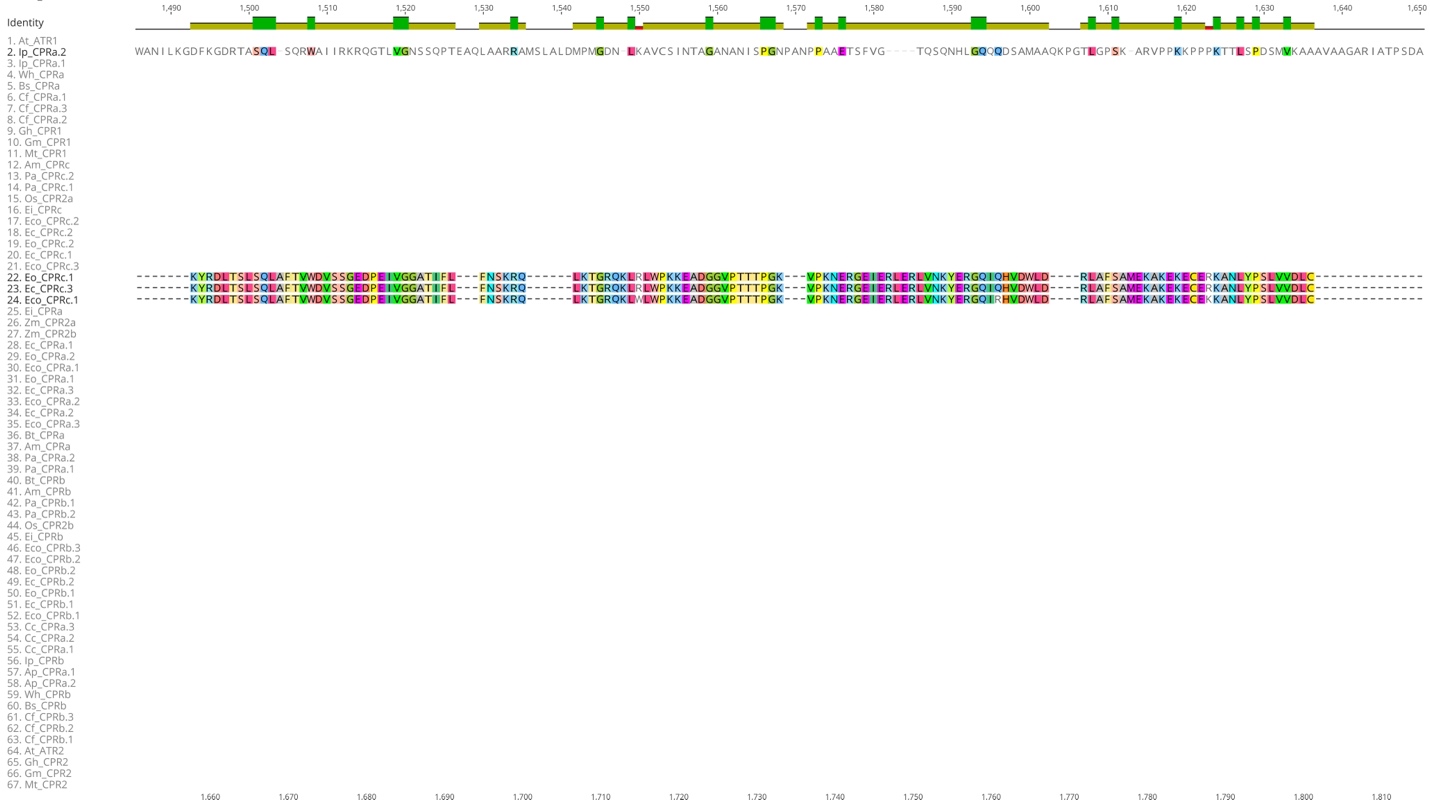

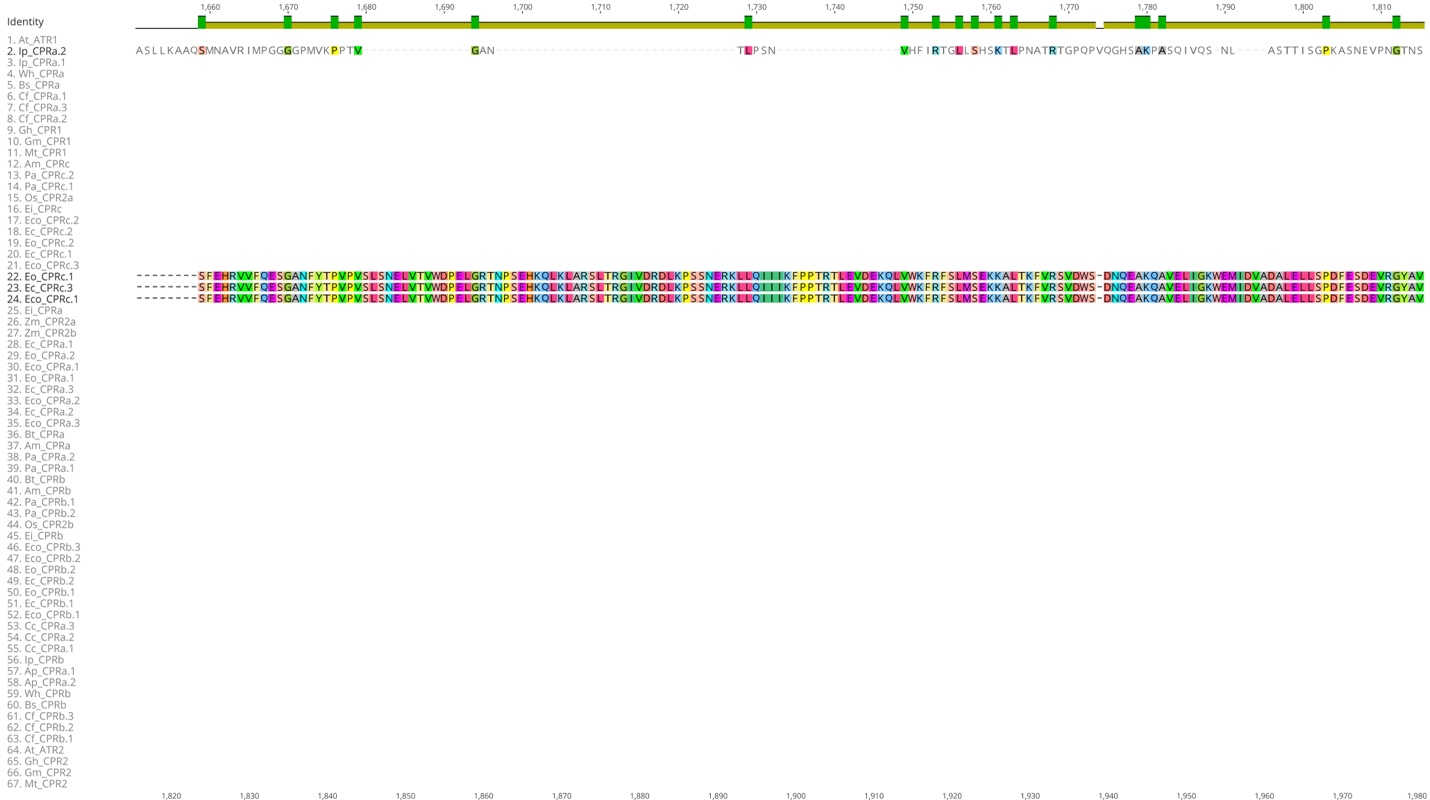

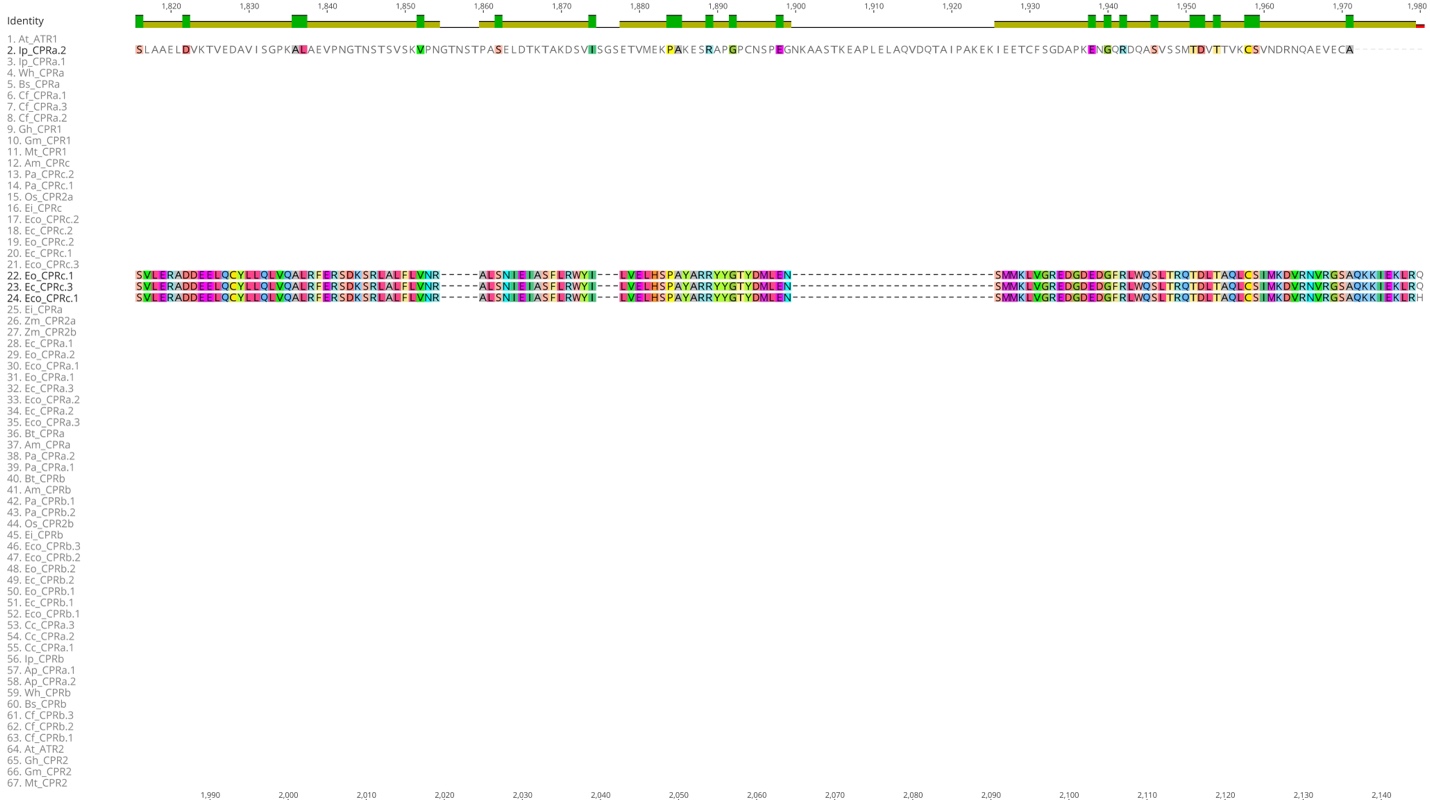

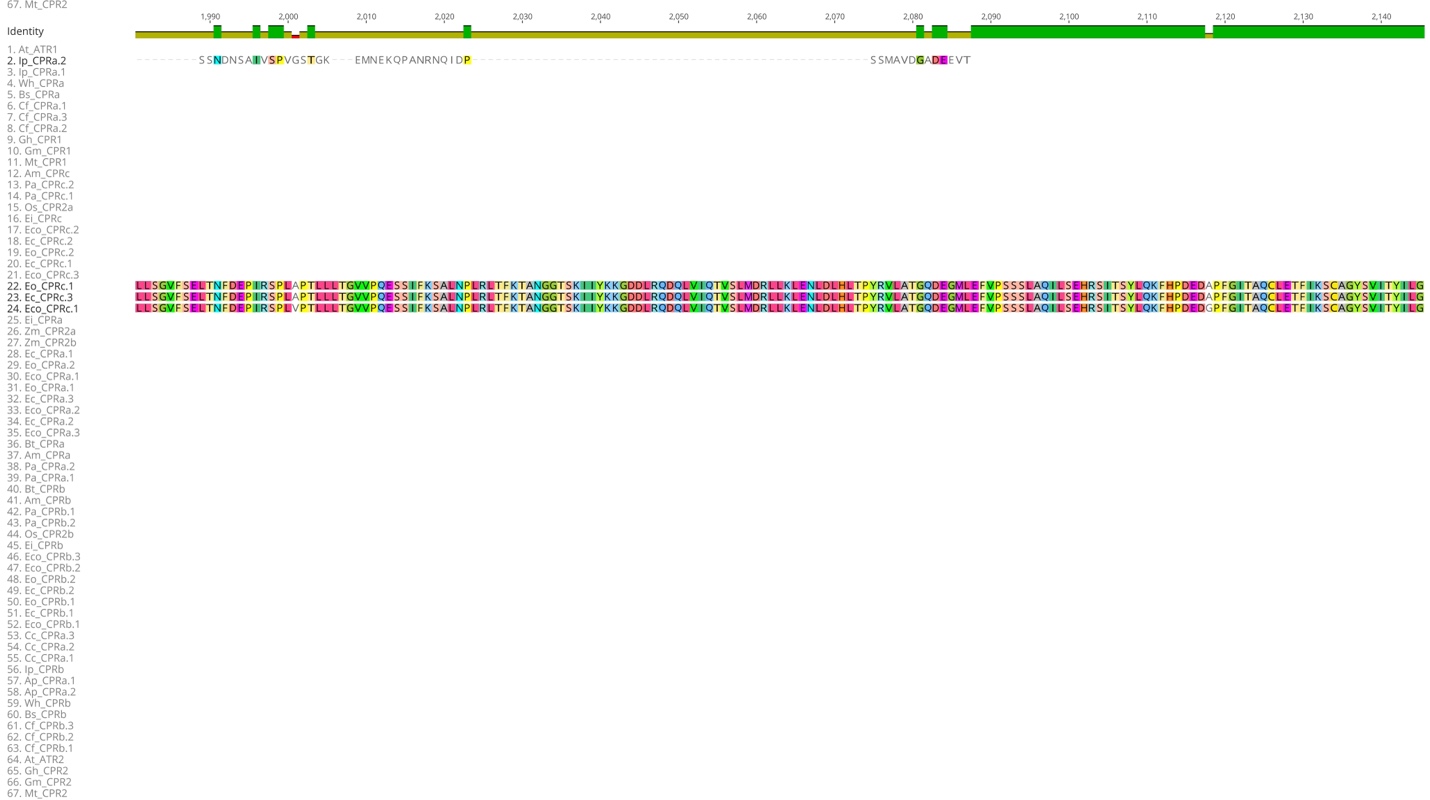

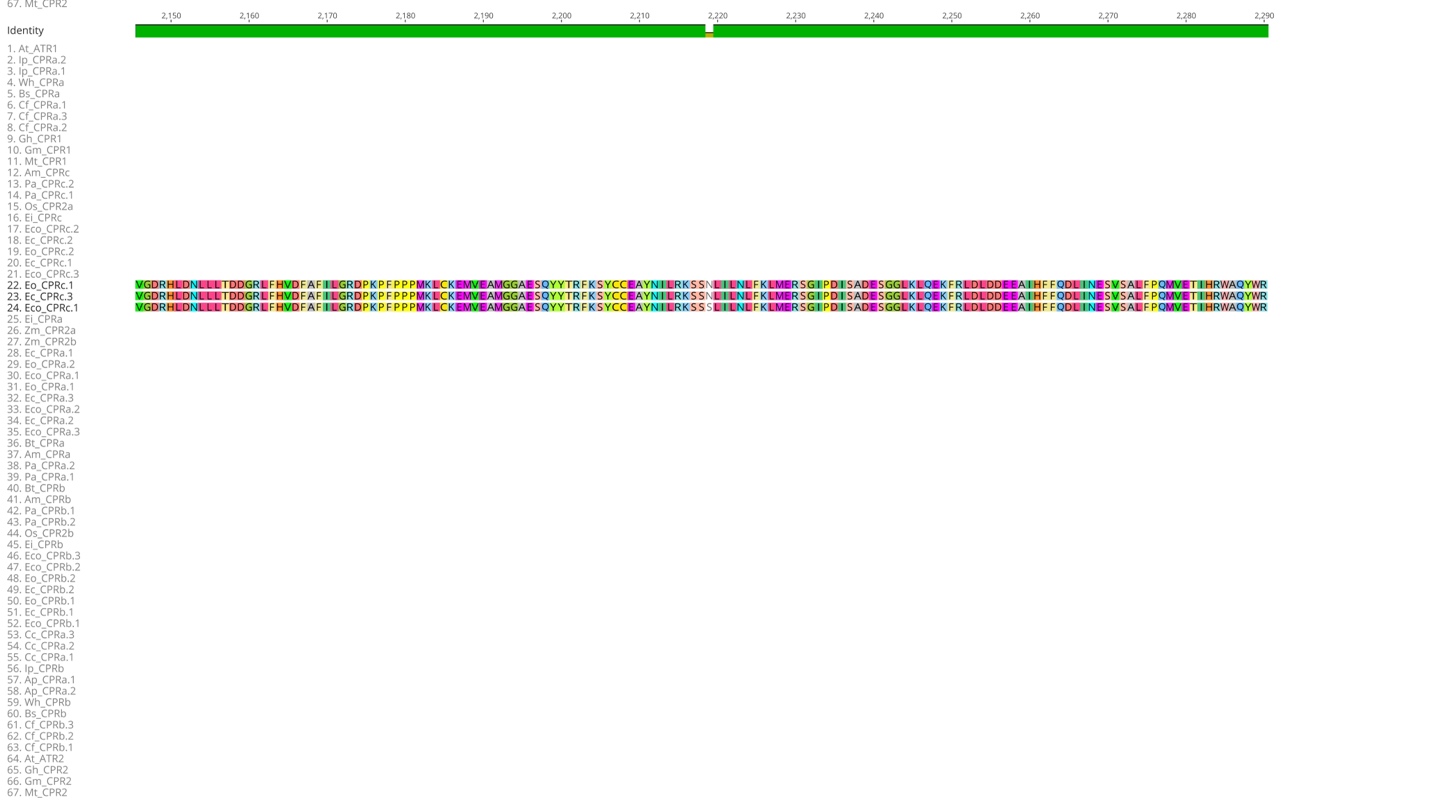

**Fig. S9.** Multiple protein alignment of 67 CPR protein sequences from 13 different weed species and others six plant species with colors indicating conserved residue between sequences. Transmembrane (347-366), FMN binding (418-665), FAD binding (727-950) and NADP binding (999-1109) are indicated. Different CPR within species were named as “a”, “b” or “c” and the copies were numbered as “.1”, “.2” or “.3” for one, two or three copies, respectively. Am - *Alopecurus myosuroides* (blackgrass), At - *Arabidopsis thaliana* (Arabidopsis), Ap – *Amaranthus palmeri* (Palmer amaranth), Wh – *Amaranthus tuberculatus* (waterhemp), Bs - *Bassia scoparia* (Kochia), Bt - *Bromus tectorum* (downy brome), Cc - *Conyza canadensis* (horseweed), Cf - *Chenopodium formosanum* (rainbow rice), Ec - *Echinochloa crus-galli* (barnyardgrass)*, Eo - E. oryzicola* (late watergrass), Eco - *E. colona* (jungle rice), Ei - *Eleusine indica* (goosegrass), Gh - *Gossypium hirsutum* (cotton), Gm - *Glycine max* (soybean), Ip - *Ipomoea purpurea* (common morning glory), Mt - *Medicago truncatula* (Barrel medic), Os - *Oryza sativa* (rice), Pa - *Poa annua* (annual bluegrass). Note: the sequence 22, 23 and 24, which correspond to Eo_CPRc.1, Ec_CPRc.3 and Eco_CPRc.1, respectively, have a C-term extensions matching phosphatidylinositol 3-kinase, which could be an assembly error.

**Table S1.** Primers and their characteristics used to genotype by PCR *ATR1* and *ATR2* genes with the T-DNA insertion and *CYP81A12* in F1 or F2 plants.

| **Gene** | **line** | **primer** | **sequence 5' to 3'** | **amplicon size (bp)** |
| --- | --- | --- | --- | --- |
| *ATR1* (AT4G24520) | *atr1-a* | LP | ATCATCGGCAGCATAGTCATC | 1045 |
|  |  | RP | AATGAATGGGCACAAAAACAG |  |
|  | *atr1-b* | LP | TGATTCTTTCCTGCAAACCAC | 1098 |
|  |  | RP | TTATCCGAAGAAATCAAAGCG |  |
| *ATR2* (AT4G30210) | *atr2-a* | LP | ACCTGAACAACCCACTCAATG | 1085 |
|  |  | RP | ATTGGTTGCATCGTTATGCTC |  |
|  | *atr2-b* | LP | TCACTCTTCTCGTAAGGCAC | 1173 |
|  |  | RP | GCTTCATACTCCCGAGTCTG |  |
| - | - | LBb1.3 | ATTTTGCCGATTTCGGAAC |  |
| *CYP81A12* | - | *Forward* | ATGGATAAGGCCTACGTGGCC | 1557 |
|  | - | *Reverse* | TCAGAGCTCCTGAAGAACATCATG |  |

LP – left border, RP – right border, LBb1.3 - primer designed in the right border of the T-DNA sequence.

**Table S2.** Primers and their characteristics used for gene expression analysis of *ATR1*, *ATR2*, and CYP81A12.

| Gene | Exp.^1^ | primer | Comment | sequence 5' to 3' | size (bp) |
| --- | --- | --- | --- | --- | --- |
| *ATR1* | qPCR | ATR1_A | *Forward* (spanning the *atr1-a* T-DNA insertion site) | CAAATCGGAAGCATACAAAG | 167 |
|  |  |  | *Reverse* (spanning the *atr1-a* T-DNA insertion site) | CACCACCGATAACTCAATTA |  |
|  |  | ATR1_B | *Forward* | ATCATCCTCGTCCTTAGCCATA | 172 |
|  |  |  | *Reverse* | TATGGGGACAGATTCGTTATCC |  |
|  |  | ATR1_C | *Forward* (spanning the *atr1-b* T-DNA insertion site) | TGTTCATATTGGCGATTACC | 130 |
|  |  |  | *Reverse* (spanning the *atr1-b* T-DNA insertion site) | CCTACTGACAATGCTGCCA |  |
|  |  | ATR1_D | *Forward* | TCCTTCTGCACAGCAACATC | 103 |
|  |  |  | *Reverse* | CTGAATACCGGGTGGTGACT |  |
| *ATR2* | qPCR | ATR2_A | *Forward* | GTCCGTAGCTGCTGAATTATCC | 103 |
|  |  |  | *Reverse* | GAGCATAACGATGCAACCAATA |  |
|  |  | ATR2_B | *Forward* (spanning the *atr2-a* T-DNA insertion site) | GAAGAAAGAGGATGTGGCTT | 85 |
|  |  |  | *Reverse* (spanning the *atr2-a* T-DNA insertion site) | CCATTTGTAGAATCTCGCTGC |  |
|  |  | ATR2_C | *Forward* | TGGAATTTGACATTGCTGGA | 167 |
|  |  |  | *Reverse* | GATTGGTGTGCCGTCTTCTT |  |
|  |  | ATR2_D | *Forward* (spanning the *atr2-b* T-DNA insertion site) | GTCAAAGAAGTCTACTTGAGGT | 171 |
|  |  |  | *Reverse* (spanning the *atr2-b* T-DNA insertion site) | TCTCATAAACCAGTGCACATG |  |
|  |  | ATR2_E | *Forward* | TCCAGGGACTGGATTAGCTC | 93 |
|  |  |  | *Reverse* | AACAAAACTGATGGCCCAAG |  |
| *ALS* | qPCR and ddPCR | ALS_A | *Forward* | CATATGCTTGGAATGCATGG | 99 |
|  |  |  | *Reverse* | CGTGACACGATCATCAAACC |  |
| *CYP81A12* | qPCR and ddPCR | CYP | *Forward* | ACCTCCAGAGCATCATCCAC | 136 |
|  |  |  | *Reverse* | TACACGTTCACCAGCAGCAT |  |

^1^ Experiment, qPCR – primers were used for real-time qPCR gene expression analyses for the respective gene. ddPCR, primer used for genotyping the copy number of the respective gene by digital droplet PCR.

**Table S3.** PCR genotyping for T-DNA insertion in F2 segregating plants derived from crosses between *atr1* or *atr2* mutants with transgenic Arabidopsis *CYP81A12*.

| Cross | Gene | Genotype | plants | frequency (%) |
| --- | --- | --- | --- | --- |
| A | *ATR1* | HM-WT | 7 | 12.1 |
|  |  | HM-M | 15 | 25.8 |
|  |  | HZ | 36 | 62.1 |
|  |  | total | 58 |  |
| B | *ATR1* | HM-WT | 16 | 25.8 |
|  |  | HM-M | 12 | 19.3 |
|  |  | HZ | 34 | 54.8 |
|  |  | total | 62 |  |
| C | *ATR2* | HM-WT | 20 | 34.8 |
|  |  | HM-M | 9 | 15.5 |
|  |  | HZ | 29 | 50 |
|  |  | total | 58 |  |
| D | *ATR2* | HM-WT | 9 | 17.3 |
|  |  | HM-M | 15 | 28.8 |
|  |  | HZ | 28 | 53.84 |
|  |  | total | 52 |  |

HM-WT – homozygous wild type, HM-M – homozygous mutant, HZ – heterozygous. Cross A – transgenic *CYP81A12* 🞨 *atr1-a*, cross B – transgenic *CYP81A12* 🞨 *atr1-b*, cross C – transgenic *CYP81A12* 🞨 *atr2-a*, and cross D – transgenic *CYP81A12* 🞨 *atr2-b*.

**Table S4.** ddPCR results of parental transgenic Arabidopsis *CYP81A12* and four selected F3 plants from crosses A (*CYP81A12* 🞨 *atr1-a*), B (*CYP81A12* 🞨 *atr1-b*), C (*CYP81A12* 🞨 *atr2-a*), and D (*CYP81A12* 🞨 *atr2-b*).

| Cross | Samples | conc (copies μL^-1^) | | | | | | ALS/  CYP81 |
| --- | --- | --- | --- | --- | --- | --- | --- | --- |
|  |  | *ALS* | Pmin^1^ | Pmax^2^ | *CYP81A12* | Pmin | Pmax |  |
| Parental | *CYP81A12* | 25.2 | 21.5 | 26.4 | 24 | 22.6 | 27.9 | 1.1 |
|  | *CYP81A12* | 12.6 | 11.7 | 15.4 | 13.6 | 10.8 | 14.4 | 0.9 |
| A | *CYP81 atr1-a* | 86 | 81 | 92 | 77.6 | 72.7 | 82.5 | 1.1 |
|  | *CYP81 atr1-a* | 110 | 104 | 116 | 104 | 99 | 110 | 1.1 |
|  | *CYP81 atr1-a* | 135 | 128 | 141 | 120 | 114 | 126 | 1.1 |
|  | *CYP81 atr1-a* | 116 | 110 | 122 | 109 | 103 | 115 | 1.1 |
| B | *CYP81 atr1-b* | 72.7 | 68.1 | 77.4 | 72.1 | 67.4 | 76.8 | 1.0 |
|  | *CYP81 atr1-b* | 83.1 | 78.1 | 88 | 74 | 69 | 79 | 1.1 |
|  | *CYP81 atr1-b* | 50.8 | 47 | 54.7 | 49.7 | 45.7 | 53.6 | 1.0 |
|  | *CYP81 atr1-b* | 78.6 | 73.8 | 83.4 | 75 | 70 | 80 | 1.0 |
| C | *CYP81 atr2-a* | 187 | 194 | 208 | 200 | 179 | 194 | 0.9 |
|  | *CYP81 atr2-a* | 115 | 109 | 120 | 111 | 105 | 117 | 1.0 |
|  | *CYP81 atr2-a* | 67.2 | 62.8 | 71.6 | 59.8 | 55.4 | 64.2 | 1.1 |
|  | *CYP81 atr2-a* | 178 | 170 | 185 | 176 | 168 | 183 | 1.0 |
| D | *CYP81 atr2-b* | 45.1 | 41.3 | 48.8 | 45.7 | 41.8 | 49.6 | 1.0 |
|  | *CYP81 atr2-b* | 86 | 81 | 91 | 89 | 84 | 94 | 1.0 |
|  | *CYP81 atr2-b* | 77 | 72 | 82 | 75.2 | 70.4 | 80.2 | 1.0 |
|  | *CYP81 atr2-b* | 98 | 93 | 104 | 89 | 84 | 94 | 1.1 |

^1^Pmin – Minimum target concentration normalized for the low error bar of the droplet Poisson distribution for the 95% confidence interval. ^2^Pmax – Maximum target concentration normalized for the high error bar of the droplet Poisson distribution for the 95% Confidence Interval.

**Table S5.** Genotype analysis of F2 segregating populations derived from crosses between *atr1* and *atr2* Arabidopsis mutants.

| Crosses | F1 plant | ATR1> | HZ | HM-WT | HZ | HZ | HM-M | HM-WT | HM-WT | HM-  M | HM-  M | Total |
| --- | --- | --- | --- | --- | --- | --- | --- | --- | --- | --- | --- | --- |
|  |  | ATR2> | HZ | HZ | HM-WT | HM-M | HZ | HM-WT | HM-M | HM-WT | HM-  M |  |
| *atr1-a* 🞨 *atr2-a* | F1  plant I | Plants | 7 | 2 | 2 | 3 | 0 | 0 | 9 | 14 | 0 | 37 |
|  |  | (%) | 18.9 | 5.4 | 5.4 | 8.1 | 0.0 | 0.0 | 24.3 | 37.8 | 0.0 | 100 |
| *atr1-a* 🞨 *atr2-a* | F1  plant II | Plants | 16 | 1 | 2 | 4 | 1 | 0 | 8 | 3 | 0 | 35 |
|  |  | (%) | 45.7 | 2.9 | 5.7 | 11.4 | 2.9 | 0.0 | 22.9 | 8.6 | 0.0 | 100 |
| *atr1-a* 🞨 *atr2-a* | F1  plant III | Plants | 27 | 1 | 2 | 3 | 0 | 0 | 12 | 11 | 0 | 56 |
|  |  | (%) | 48.2 | 1.8 | 3.6 | 5.4 | 0.0 | 0.0 | 21.4 | 19.6 | 0.0 | 100 |
| *atr1-a* 🞨 *atr2-a* | F1  plant IV | Plants | 16 | 0 | 0 | 3 | 1 | 0 | 7 | 9 | 0 | 36 |
|  |  | (%) | 44.4 | 0.0 | 0.0 | 8.3 | 2.8 | 0.0 | 19.4 | 25.0 | 0.0 | 100 |
| *atr1-b* 🞨 *atr2-a* | F1  plant I | Plants | 20 | 1 | 2 | 1 | 1 | 0 | 16 | 6 | 0 | 47 |
|  |  | (%) | 42.6 | 2.1 | 4.3 | 2.1 | 2.1 | 0.0 | 34.0 | 12.8 | 0.0 | 100 |
| *atr1-b* 🞨 *atr2-a* | F1  plant II | Plants | 42 | 2 | 2 | 5 | 2 | 0 | 19 | 14 | 0 | 86 |
|  |  | (%) | 48.8 | 2.3 | 2.3 | 5.8 | 2.3 | 0.0 | 22.1 | 16.3 | 0.0 | 100 |
| *atr1-b* 🞨 *atr2-b* | F1  plant I | Plants | 64 | 1 | 3 | 1 | 1 | 0 | 27 | 22 | 0 | 119 |
|  |  | (%) | 53.8 | 0.8 | 2.5 | 0.8 | 0.8 | 0.0 | 22.7 | 18.5 | 0.0 | 100 |
| Total | | Plants | 192 | 8 | 13 | 20 | 6 | 0 | 98 | 79 | 0 | 416 |
|  |  | (%) | 46.2 | 1.9 | 3.1 | 4.8 | 1.4 | 0.0 | 23.6 | 19.0 | 0.0 | 100 |
| Expected | | (%) | 25 | 12.5 | 12.5 | 12.5 | 12.5 | 6.25 | 6.25 | 6.25 | 6.25 | 100 |
| χ^2^ test | | χ^2^ = 560.75 | | | df = 8 | | | | p-value = <0.0001 | | | |

HM-WT, homozygous wildtype, HM-MT, homozygous mutant, HZ – heterozygous. *ATR1* – Arabidopsis P450 reductase 1, *ATR2* - Arabidopsis P450 reductase 2.

**Table S6.** Weed species and their information about NADPH-cytochrome P450 reductase composition.

| species | common name | ploidy | #chr | #CPR | # copies | Gene ID | CPR class | source |
| --- | --- | --- | --- | --- | --- | --- | --- | --- |
| *Alopecurus myosuroides* | blackgrass | 2n | 7 | 3 | 1 | ALOMY3G10997  ALOMY7G41785  ALOMY6G43414 | II | (Cai *et al.*, 2023) |
| *Amaranthus*  *palmeri* | Palmer amaranth | 2n | 34 | 3 | 1 | AmaPaChr07Bg124140  AmaPaChr08Bg138360  AmaPaChr14Bg215950 | I and II | (CropPedia, 2024) |
| *Amaranthus tuberculatus* | waterhemp | 2n | 32 | 3 | 1 | AmaTu_altChr06g109800  AmaTu_altChr04g073760  AmaTu_altChr12g193200 | I and II | (CropPedia, 2024) |
| *Bassia scoparia* | kochia | 2n | 9 | 2 | 1 | Bs.00g055750  Bs.00g138840 | I and II | (Hall *et al.*, 2023) |
| *Bromus tectorum* | downy brome | 2n | 7 | 2 | 1 | Bt030022  Bt011718 | II | (Revolinski *et al.*, 2023) |
| *Chenopodium formosanum* | Djulis | 6n | 27 | 2 | 3 | CheFoCf8Cg733520  CheFoCf1Bg010510  CheFoCf1Dg073860  CheFoCf2Cg136610  CheFoCf8Bg696010  CheFoCf8Dg788700 | I and II | (Jarvis *et al.*, 2022) |
| *Conyza canadensis* | horseweed | 2n | 9 | 3 | 1 | g12026  g21967  g20812 | II | (Laforest *et al.*, 2020) |
| *Echinochloa colona* | jungle rice | 6n | 27 | 3 | 3 | FL09.2481  EL09.2594  DL09.2856  EL03.2838  FL03.2645  DL03.3340  EL08.707  DL08.911  FL08.729 | II | (Wu *et al.*, 2022) |
| *Echinochloa crus-galli* | barnyardgrass | 6n | 27 | 3 | 2 and 3 | AH09.2541  CH09.2973  BH09.2775  CH03.3272  BH03.3081  CH08.947  AH08.811  AH07.1112 | II | (Wu *et al.*, 2022) |
| *Echinochloa oryzicola* | late watergrass | 4n | 18 | 3 | 2 | BT09.2442  AT09.2615  BT03.3019  AT03.3018  BT08.1100  AT08.836 | II | (Wu *et al.*, 2022) |
| *Eleusine indica* | goosegrass | 2n | 9 | 3 | 1 | Eindica.09G039898  Eindica.03G015096  Eindica.06G031355 | II | (Zhang *et al.*, 2023) |
| *Ipomoea purpurea* | common morning-glory | 2n | 15 | 2 | 1 and 2 | Ipurp_gene3295  Ipurp_gene24259  Ipurp_gene29825 | I and II | (Gupta *et al.*, 2023) |
| *Poa annua* | annual bluegrass | 4n | 14 | 3 | 2 | PoaAnPa1Bg125530  PoaAnPa1Ag065050  PoaAnPa7Ag697440  PoaAnPa5Ag490280  PoaAnPa7Bg739430  PoaAnPa5Bg534380 | II | (Robbins *et al.*, 2023) |

#chr – species chromosome number, #CPR – number of NADPH cytochrome P450 reductase.

**Methods S1** Characterization and confirmation of T-DNA insertion

PCR assays were performed to identify and confirm the presence of T-DNA insertion in the *ATR1* and *ATR2* genes in the mutant lines and to verify homozygosity. To amplify the wild type (WT) version of *ATR1* or *ATR2* genes, different primer sets named LP (forward) and RP (reverse) spanning the T-DNA insertion specific for each gene and line were used. The primers to identify the T-DNA insertion for all the mutant lines were RP primers, specific for each gene and line, with LBb1.3 (Fig. **S1**). The primer LBb1.3 is a primer designed in the right border of the T-DNA sequence. Primer sequences of LBb1.3, LP and RP for *atr1-a*, *atr1-b* and *atr2-a* were provided by SIGnAL (http://signal.salk.edu/). Primer sets for *atr2-b* were designed using Primer3plus. The primer sequences are described in Tab. **S1**.

Leaf tissue was collected from individual plants at rosette stage and incubated in 1.2 mL tubes. Around 20 plants for each mutant line were genotyped. The tissues were frozen at -20°C until further analyses. Frozen materials were ground using a Qiagen Tissuelyzer and DNA was extracted using a modified hexadecyltrimethylammonium bromide (CTAB) method. DNA quality and concentration were measured using Nanodrop2000.

PCR reactions were performed with Econotaq Plus 2🞨 Master Mix. PCR mixtures consisted of 12.5 µL of Econotaq Master Mix, 6.5 µL nuclease-free water, and 2.5 µL of forward and reverse primers (10 µM). Each sample was submitted to two different reactions, called “WT PCR” and “T-DNA PCR”. The former consisted of LP and RP primers to amplify the WT form or heterozygous and the latter was composed of RP and LBb1.3 primers to amplify the T-DNA insertion, if present. PCR cycling conditions were composed by initial denaturation of 94°C for 2 min, followed by 35 cycles consisting of denaturation at 94°C for 30 s, annealing at 55°C for “WT PCR” and 48°C for “T-DNA PCR” for 30 s, followed by extension at 72°C for 70 s, with a final extension time at 72°C for 5 min. The DNA amplification was confirmed by running in 1% agarose gel electrophoresis and PCR products were sent for Sanger sequencing.

**Methods S2** Generating transgenic herbicide-resistant Arabidopsis with mutant *atr1* or *atr2* genes

Transgenic Arabidopsis *CYP81A12* and mutant lines *atr1-a*, *atr1-b*, *atr2-a*, and *atr2-b* were grown with the same conditions as indicated before. When plants were in the flowering stage, the mutant lines were crossed with the transgenic Arabidopsis *CYP81A12*. The cross was repeated three additional times with new flowers within an interval of a week. The crossed flowers were labeled, and the seeds were collected when siliques turned brown. In total, four crosses were performed, cross A – transgenic *CYP81A12* 🞨 *atr1-a*, cross B – transgenic *CYP81A12* 🞨*atr1-b*, cross C – transgenic *CYP81A12* 🞨*atr2-a*, and cross D – transgenic *CYP81A12* 🞨 *atr2-b* (Fig. **2**).

F1 plants were grown, and tissue collected two weeks after transplanting for DNA extraction. Extracted DNA was used to genotype for T-DNA insertion and presence of *CYP81A12* gene. T-DNA insertion was genotyped as indicated before, using WT PCR and T-DNA PCR reactions. PCR amplification for the whole *CYP81A12* was performed to genotype for the presence of the gene using genomic DNA. Primer sequences for *CYP81A12* are listed in Tab. **1**. Plants that were heterozygous for T-DNA insertion and had the presence of *CYP81A12* were selected and self-pollinated. F2 plants were grown with the same conditions as described before, and tissue collected for genotyping. Genomic DNA was extracted, and T-DNA insertion was genotyped using the same two PCR reactions as indicated before. It was expected that 25% of the F2 population were homozygous for T-DNA insertion. These plants were selected to genotype for the presence of *CYP81A12* and those plants that had the gene were selected. These plants were genotyped for copy number analysis of *CYP81A12* using QX200 Droplet Digital PCR system (ddPCR) due to its precision and sensitivity for detection of low copy numbers. ddPCR consisted of a master mix composed of 10 µL of 2🞨QX200 EvaGreen^®^ Supermix, 2.5 µL of each primer (5 µM), and 5 µL of genomic DNA (5 ng µL^-1^).

Selected samples were processed for droplet generation using QX200™ droplet generator, followed by PCR reaction with initial cycle of 5 min at 95°C followed by a sequence of 40 cycles starting at 95°C for 30 s and 60°C for 1 min. Droplet reading of PCR products was conducted in the QX200 Droplet Reader. The *ALS* gene was used as a reference gene containing only two copies (one copy per haplotype). Primer sequences for *ALS* and *CYP81A12* used for ddPCR are listed in Tab. **S2**. The samples were analyzed by QuantaSoft AP Software, which measures the number of positive and negative droplets for each gene in each well. Droplets are assigned as positive or negative by thresholding based on their fluorescence amplitude. The software calculates the starting concentration of each target DNA molecule by modeling a Poisson distribution; the formula used for Poisson modeling is: Copies per droplet = -ln(1-p) where p= fraction of positive droplets. Concentrations are provided in units of copies per microliter of input sample with 95% confidence intervals. The ratio of the number of copies per droplet of *ALS* and *CYP81A12* was calculated.

**Methods S3** Generating double knockout *atr1* *atr2* Arabidopsis lines

To obtain an Arabidopsis line with a double knockout of *ATR1* and *ATR2*, mutant lines *atr1-a*, *atr1-b*, *atr2-a*, and *atr2-b* were cultivated under the same conditions as previously indicated. Upon reaching the flowering stage, crosses were performed as described earlier. F1 plants were subjected to genotyping to determine homozygosity for *ATR1* and *ATR2* using the same PCR reactions as previously mentioned. Heterozygous plants were selected for self-pollination. F2 seeds were then sown in soil, and subsequent plants were genotyped for *atr1* and *atr2* mutations.
